# Interplay between purging and admixture shapes genetic load in an invasive guppy population

**DOI:** 10.1101/2025.09.12.675788

**Authors:** Katarzyna Burda, Mary J Janecka, Ryan S. Mohammed, David R. Clark, Rachael Kramp, Jessica F. Stephenson, Jacek Radwan, Mateusz Konczal

## Abstract

Demographic history can shape the genetic load of populations by influencing the efficacy of selection, levels of heterozygosity, and the incorporation of new variants via gene flow. Understanding these dynamics is crucial for identifying threats to population viability and predicting evolutionary trajectories of invasive populations translocated by humans into non-native environments. We investigate these processes in Trinidadian guppies (*Poecilia reticulata*) by estimating genetic loads across multiple populations, with a particular focus on a single expanding population in which translocated individuals have rapidly spread and replaced native individuals. Overall, we observe the expected negative relationship between neutral genetic diversity and relative genetic load. In the translocated population, patterns differ between strongly and weakly deleterious mutations. Strongly deleterious alleles are purged at the isolated translocation site but tend to accumulate along the expansion front. In contrast, the genetic load estimated based on weakly deleterious variants declines along the expansion gradient. These differing patterns can be explained by admixture with native populations (which carried fewer weakly deleterious mutations) reducing the overall genetic load of the population at the expansion front. However, admixture has also increased genetic diversity and introduced new strongly deleterious alleles, thereby reversing the purging effect. Together, our findings illustrate the complex interactions determining genetic load in subdivided populations, offering important insights into the evolutionary aspects of biological invasions.

## INTRODUCTION

Genetic load refers to the decrease in fitness caused by deleterious variants in a genome (Crow 1989; Bertorelle et al. 2022). Most of the deleterious mutations have slight effects in heterozygous carriers and remain rare in large panmictic populations (Eyre-Walker and Keightley 2007). Such deleterious variants present in heterozygous genotypes are often referred to as “heterozygous genetic load” (Bertorelle et al. 2022). Under some demographic scenarios, deleterious variants can increase in frequency and become exposed in homozygotes as a “homozygous genetic load”, which may threaten population survival (Mathur and DeWoody 2021). In such a situation, deleterious alleles may either become common, or even fixed, in a population decreasing its mean fitness, or they may be eliminated from the population by “purging” (Dussex et al. 2023; Robinson et al. 2023).

Demographic changes, such as a population reduction, are expected to affect genetic variation and the genetic load of a population through increased genetic drift (Wright 1932) and inbreeding (Wright 1922). Genetic drift increases the variance around the mean frequency of deleterious mutations, allowing some deleterious alleles to drift to high frequencies (de Pedro et al. 2021; Zeitler et al. 2023; Stewart et al. 2017). Thus, in small populations in which selection fails to remove deleterious mutations of weak effects, these variants accumulate over time (Kimura and Ohta 1969) and may eventually fix (Kimura et al. 1963). This accumulation reduces fitness, which may further reduce population size, potentially resulting in a vicious circle of “mutational meltdown” (Lynch et al. 1995). Under this scenario, such “genetic erosion” in small, isolated populations of endangered species should relate to their estimated neutral genetic variation, reflecting “effective population size”. In other words, genome-wide genetic variation underlies fitness and thus provide crucial information for conservation (Kardos et al. 2021; DeWoody et al. 2021; Willi et al. 2022). Consistent with these theoretical predictions, a negative relationship between neutral nucleotide diversity and homozygous load was indeed found, for example in an endangered rattlesnake (Mathur et al. 2023a) or Montezuma quail (Mathur et al. 2023b).

However, the relationship between genetic load and neutral diversity depends on the timescale of evolutionary processes that generate these correlations (Mathur et al. 2023a, 2023b) and may be changed by recent demographic events. For instance, a recent transient population size reduction might increase a population’s mean fitness through several processes. First, alleles at low frequencies, such as strongly deleterious mutations, are likely to be lost by drift. Second, reduced population size increases the probability that alleles carried by mating partners are identical by descent, leading to increased homozygosity in a population (Wright 1922) thus changing masked load into realized load. Highly deleterious variants at low frequencies in the pre-bottleneck population thus become exposed to natural selection (Charlesworth and Willis 2009). If selection can overcome genetic drift, it may purge large effect deleterious variants, ultimately decreasing both heterozygous and total genetic load (Glémin 2003; Crnokrak and Barrett 2002; Keller and Waller 2002; Barrett and Charlesworth 1991). New genomic data suggest that purging occurs more often than previously thought (Dussex et al. 2023), however, its effectiveness differs between species (Lan et al. 2025) and its consequences for population viability and extinction risk remain unclear (Kardos et al. 2025). Consequently, Teixeira and Huber (2021) argued against the importance of neutral genetic diversity for the conservation of wild populations, and instead advocated for a better understanding of the properties of non-neutral genetic diversity. Quantitative assessment of functional variation is, however, difficult, and despite wide discussion and studies in multiple systems, there is still no general understanding of how useful such information is for wild populations conservation (Kardos et al. 2023; van Oosterhout et al. 2026; Ralls et al. 2020; Kardos et al. 2021; Robinson et al. 2019; Kyriazis et al. 2021; Kardos et al. 2025).

Such investigations might also be important in the context of biological invasions (Estoup et al. 2016). Even though invasions and endangered species reflect opposite ends of a spectrum of ecological success, they experience similar challenges (Colautti et al. 2017). Both natural and artificial translocations that give rise to invasions usually introduce small numbers of often related individuals, and thus the initial stages of invasion involve small effective population sizes. Maintaining invasive potential despite expected loss of genetic diversity and increased genetic load was termed the “genetic paradox” of invasions (Estoup et al. 2016; Kołodziejczyk et al. 2025). It has been suggested that purging may explain this paradox and contribute to the invasive potential of translocated populations (Estoup et al. 2016). Purging in invasive species has, however, been documented in only a few cases (Facon et al. 2011; Lombaert et al. 2025; Zayed et al. 2007), making generalization difficult. Following an initial bottleneck, invasive species often experience rapid range expansion. Such expansions can involve multiple subsequent bottlenecks, increasing the effects of purging and genetic drift especially at the expansion front (Peischl et al. 2015). This process might then lead to further accumulation of weakly deleterious variants and purging of strongly deleterious ones (Rougemont et al. 2023).

However, following the initial bottleneck, successful invasions are often associated with admixture between divergent genotypes (when translocations occur multiple times) or with hybridization between translocated and native populations (Barker et al. 2019; Kołodziejczyk et al. 2025; Colautti et al. 2017). Potential benefits of such admixture include increased genetic variation and incorporation of alleles associated with local adaption into the genetic pool of invasive species, enabling adaptation to novel habitats (Wagner et al. 2017; Kołodziejczyk et al. 2025). Interestingly, genome-wide genetic diversity was not significantly reduced in many invasive populations, likely due to recent admixture or hybridization (Kołodziejczyk et al. 2025). However, little is known about the effect of these admixture events on the dynamics of genetic load. On one hand, admixture might be expected to mask the homozygous genetic load of an expanding population, not only for deleterious mutations that increased in frequency during past bottlenecks, but also for variants already fixed in a population. On the other hand, it has the potential to introduce new variants into a population, some of them being deleterious, potentially increasing the total genetic load of an invasive population. It has been suggested for example, that introgression of Neanderthal ancestry into anatomically modern human genomes, during the expansion out of Africa, reduced fitness by at least 0.5%, due to increased genetic load derived from Neanderthal genomes (Harris and Nielsen 2016). However, whether such scenarios can affect invasion potential has rarely been studied.

Here, we examine the fate of genetic load during the invasion of Trinidadian guppies (*Poecilia reticulata*) into a diverged, allopatric population, during which translocated individuals rapidly spread and replaced native individuals. Guppies occur naturally in the fresh waters of Trinidad, Tobago and South America, but have been translocated and successfully invaded multiple sites across the globe (Deacon et al. 2011). Natural Trinidadian populations have been intensively studied by generations of ecologists and evolutionary biologists (Magurran 2005) and some of these studies included translocation experiments (Fitzpatrick et al. 2016; Fraser et al. 2015). One such translocation occurred in the Turure River in 1957 (Endler 1980; Reznick and Bryga 1987; Shaw et al. 1992). Fish from the lower Guanapo River (Caroni drainage) were moved to a site in the upper Turure River (Oropouche drainage) which previously lacked guppies (Magurran 2005). 200 guppies, approximately half females, were introduced (Shaw et al. 1992; Sievers et al. 2012). Importantly, the source and recipient drainages are inhabited by differentiated guppy populations (Willing et al. 2010), with long separate histories (estimated to have split 0.18-1.2 mya, based on net divergence (Fajen and Breden 1992; Whiting et al. 2021)), and evidence of partial reproductive isolation (Russell and Magurran 2006). The Oropouche fish are more diverged from Caroni than from other fish on other islands (e.g. Tobago), suggesting that they may represent distinct species (Schories et al. 2009), though evidence for barriers to gene flow is weak (Devigili et al. 2018; Magurran 2005). The differentiation documented by these studies reflects both isolation and adaptation to different local environments (Reznick et al. 1990; Bassar et al. 2010; Kemp et al. 2018). However, after translocation, genetically foreign individuals spread rapidly, and largely replaced native individuals in the lower parts of the Turure River. Furthermore, several sources of evidence demonstrated ongoing admixture from resident populations (Becher and Magurran 2000; Willing et al. 2010; Sievers et al. 2012).

We aimed to test whether the spread of Guanapo-derived genomes into the resident populations of the Turure River was associated with changes in genetic load. In particular, we test the hypothesis (H1) that the Guanapo- derived population introduced into the upstream Turure was purged of highly deleterious mutations and now has a low overall genetic load, potentially increasing invasive potential (Sievers et al. 2012). To test this hypothesis, we compared the genetic load of the upstream Turure population with that of the Guanapo source population. Additionally, we explore the accumulation of genetic load along the Turure River, to test the hypothesis (H2) of expansion associated with a series of bottlenecks, allowing the accumulation of weakly deleterious mutations and the purging of strongly deleterious mutations in downriver sites. However, if admixture occurred along with expansion, as earlier work suggests (Sievers et al. 2012; Willing et al. 2010), genetic load might also be affected by this process. We therefore explore differences in both genetic load and admixture along the river. Additionally, to provide a benchmark for interpreting genetic load in our focal populations, we test for associations between neutral diversity and genetic load across multiple populations from other rivers. Guppy populations in Trinidadian rivers range in population size from hundreds to tens of thousands of individuals (Whiting et al. 2021), providing the opportunity to test for such an association.

## RESULTS

### Population structure and nucleotide diversity

We analyzed 201 individuals from 15 populations (sampling sites) originating from nine rivers on Trinidad (Aripo, Caura, Guanapo, Lopinot, Santa Cruz, La Seiva, Oropouche, Quare, Turure), and two rivers on Tobago (Dog River and Roxborough; Fig. 1B; Supplemental Table S1). Five analyzed Trinidadian rivers belong to the Caroni drainage and the other four to the Oropouche drainage. Three rivers were sampled in more than one location: Quare (upper and lower), Oropouche (upper and lower), and Turure (upper, middle and lower). Samples were sequenced yielding a mean coverage of 15x (7x - 36x) and were mapped to the reference genome with over 97% reads aligned in all individuals sampled.

**Figure 1.**
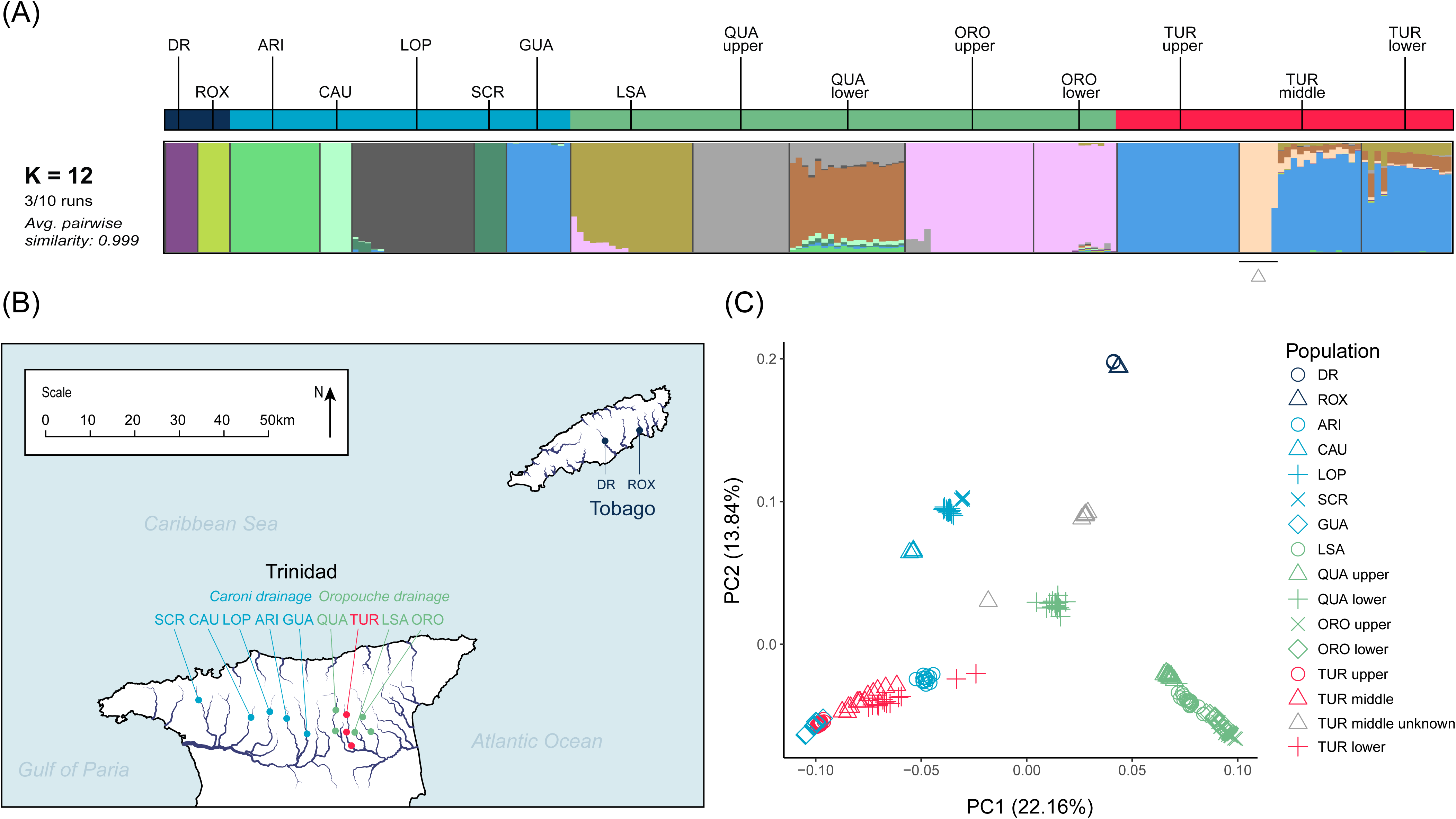
Genetic structure of the analysed dataset. **(A)** Admixture plots from analyses with optimal values of K (12). The regions are color-coded in the strip above the plots as follows: Tobago - dark blue, Caroni drainage - light blue, Oropouche drainage - green, and Turure - red. Grey triangle below the Admixture plot points to individuals of unknown origin, excluded from the analyses. (**B)** Sampling locations in Trinidad and Tobago, across the rivers: SCR (Santa Cruz), CAU (Caura), LOP (Lopinot), ARI (Aripo), Guanapo (GUA), QUA (Quare), TUR (Turure), LSA (La Seiva), ORO (Oropouche), DR (Dog River), ROX (Roxborough). **(C)** Principal Component Analysis of guppy populations based on genome-wide variation. Each point is one individual.

We performed principal component analysis (PCA) to understand the overall pattern of genetic differentiation between populations. The first PC (explaining 22.2% variation) separated Oropouche (which included the Turure, recipient river) from the Caroni (which included the Guanapo, source river) drainages, whereas the second, explained 13.8% of variation, separated samples from Trinidad from those on Tobago (Fig.1C). Turure individuals cluster with the Guanapo population. Individuals from the lower Quare River sites have intermediate positions on PC1, and cluster with neither Caroni nor other Oropuche drainage individuals, consistent with other strong signals of admixture in this population (Willing et al. 2010; Fitzpatrick et al. 2016; Whiting et al. 2021).

Admixture analysis identified the optimal K-value as 12, with clear differentiation among the two islands and the Trinidad drainages (Fig. 1A). The upper and lower Oropouche sites are highly similar to each other, whereas the upper and lower Quare sites differ, with greater variation in the lower site. In the upper Turure River, to which Caroni drainage fish were translocated, we found high similarity to the Guanapo source population and no evidence of admixture, in accordance with previous findings (Willing et al. 2010). As expected, increased admixture is detected downstream, probably reflecting crosses between introduced and native populations, although the vast majority of the ancestry is from the introduced population (Fig. 1A).

Interestingly, we observed 5 individuals from the Turure middle site, that cluster separately from the other samples from the same site (Fig. 1C). In the admixture plot they represent pure distinct genetic ancestry, while one other individual from the same population seems to be F1 hybrid, and others seem to be admixed for this ancestry (marked as grey triangles in Figs. 1A and 1C). These individuals come from two different sampling years, making it unlikely that they represent mislabeling or other errors. Most likely, these individuals represent a truly distinct genetic population within this river, possibly due to another recent translocation from an unknown source. These five individuals and the putative F1 hybrid were excluded from the analyses below.

Genome-wide nucleotide diversity estimates for 4-fold degenerated sites range from 2.3 · 10^-3^ to 6.5 · 10^-3^ per site, with the estimate for the upper Turure site being relatively similar to that for Guanapo source population (3.22 · 10^-3^ vs 2.67 · 10^-3^, Fig. 2). Analyses of the lower sites in all rivers, but excluding the post-translocation Turure populations, showed higher diversity in Oropouche (in intercept) than Caroni populations (estimate = −0.0022, se=0.0006, t= −2.899, p= 0.023) and Tobago (estimate = −0.0028, SE=0.0008, t= −3.495, p = 0.010).

**Figure 2.**
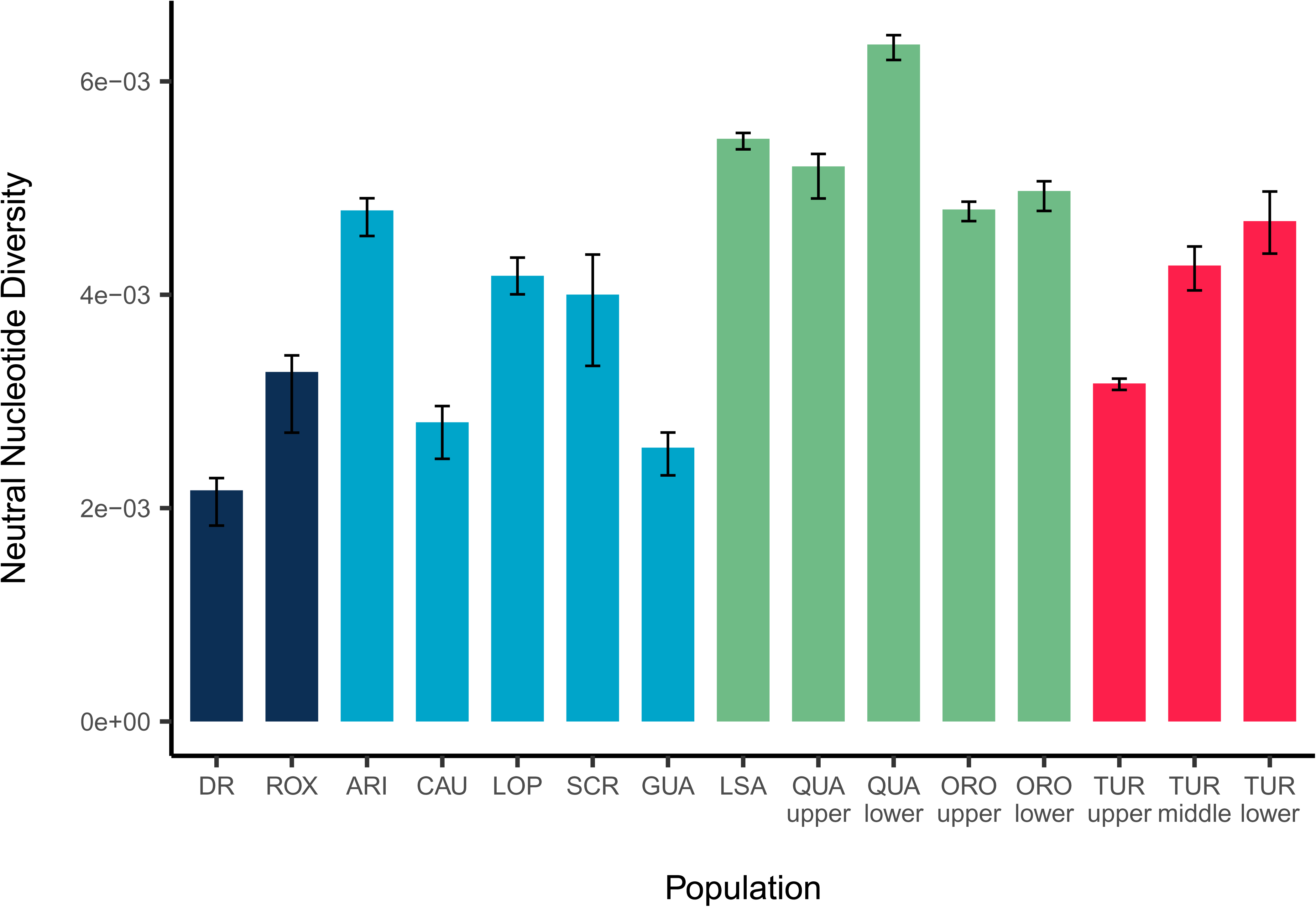
Nucleotide diversity (π) at fourfold degenerate sites across the analyzed populations. Estimates were obtained by bootstrapping individuals within each population. Bars represent the median π across 100 bootstrap replicates, and whiskers indicate the 95% bootstrap interval (2.5–97.5%). Populations are color-coded by region: Tobago (dark blue), Caroni drainage (light blue), Oropouche drainage (green), and Turure (red).

To assess whether the translocated population exhibits signals of recent bottlenecks, we estimated Tajima’s D values. Negative Tajima’s D values indicate an excess of rare variants relative to neutral expectation under drift-mutation equilibrium. This can be caused by recent population growth or by negative selection acting against mutations, whereas a positive Tajima’s D indicates an excess of common variants suggesting balancing selection, population structure, or a recent bottleneck that caused loss of rare variants. Our estimates based on 4-fold degenerated sites should thus reflect mostly demographic differences. Upstream sites showed significantly higher Tajima’s D values than downstream sites, and the upper Turure had higher values than its source Guanapo population (Supplemental Fig. S1). However, the difference between Guanapo and the upper Tururre site was smaller than that of many Oropouche populations, thus the allele frequency spectrum in this population suggests only subtle effects of the translocation event.

### Genetic load and its correlation with neutral diversity

We calculated three measures of genetic load for each re-sequenced individual: *Relative Loss-of-Function Load* (RLL), *Relative Genetic Load* (RGL) and *Relative Missense Load* (RML). The three measures are expected to reflect the effects of different classes of deleterious mutations. As loss-of-function (LoF) mutations are expected to be the most severe, with large expected effects on individual fitness on average, thus we used RLL to reflect the effects of the most deleterious class of mutations. The RGL estimate was based on conservation scores (CS) calculated from the observed number of substitutions in a multi-species alignment, relative to the number of expected substitutions under neutrality (Davydov et al. 2010). This provides information on evolutionary constraints. Mutations with high CS scores, have been used to estimate genetic load in many studies and are already a well-established approach (Bertorelle et al. 2022) that likely captures a wider range of selection coefficients than LoF mutations. Finally, many missense mutations are weakly deleterious, thus the genetic load estimated from these variants (RML) should reflect the least deleterious classes of mutations.

To estimate genetic load, the guppy reference genome was first polarized using Illumina data from two outgroup species, *Xiphophorus maculatus* and *Micropoecilia picta* (14x and 23x coverage, respectively). We inferred ancestral states for the sites used in CS analyses; 42,860,381 sites (5.9% of the genome) yielded information about both ancestral allele and CS. We called genotypes for these sites separately within each guppy population using stringent filtering criteria (see Supplemental Tab. S2 for effects of filtering on the number of called genotypes). RGL value was then calculated per individual, as the sum of presumably deleterious mutations (CS>2) weighted by the CS value, relative to the value for all derived variants. RLL and RML were respectively calculated as the ratio of loss-of-function (LoF) and missense derived variant numbers to the number of derived synonymous variants in an individual’s genome. All three values (RGL, RLL and RML) were calculated separately for homozygous and heterozygous genotypes. The sum of these two approximates the total genetic load. We used the value of CS>2 as a threshold to balance selectivity and sample size (higher CS values included very small numbers of SNPs; Supplemental Table S3). In total, we analyzed 1659383 sites, including 3479 with CS > 2. The coding regions were enriched in polymorphic sites with CS>2, when compared with intergenic regions, and nonsynonymous variants were enriched in such sites when compared with synonymous ones, demonstrating that conservation scores can capture deleterious variation. Populations show considerable variation in the mean number of putatively deleterious variants per individual despite little variation in coverage across populations (Supplemental Table S4).

Genetic load differed among geographic regions, with Oropouche populations generally having the lowest load, significantly lower than Tobago in most comparisons (total RGL, homozygous RGL, total RML, homozygous RML and heterozygous RLL, Supplemental Table S5) and lower than Caroni for total RGL (Supplemental Table S5). Consequently, in models predicting genetic load from nucleotide diversity in a population after accounting for regional differences we only found a significant negative association between nucleotide diversity and total RGL (estimate = −2.451, SE=0.798, t= −3.069, P=0.022; Fig. 3), and the region was not significant in any of these analyses. After removing the region, we found that total RGL and total RML were negatively associated with nucleotide diversity, whereas total RLL was not significantly predicted by nucleotide diversity (Supplemental Table S6, Fig. 3). Nucleotide diversity was also found to be negatively associated with homozygous RGL (Supplemental Table S6, Supplemental Fig. S2).

**Figure 3.**
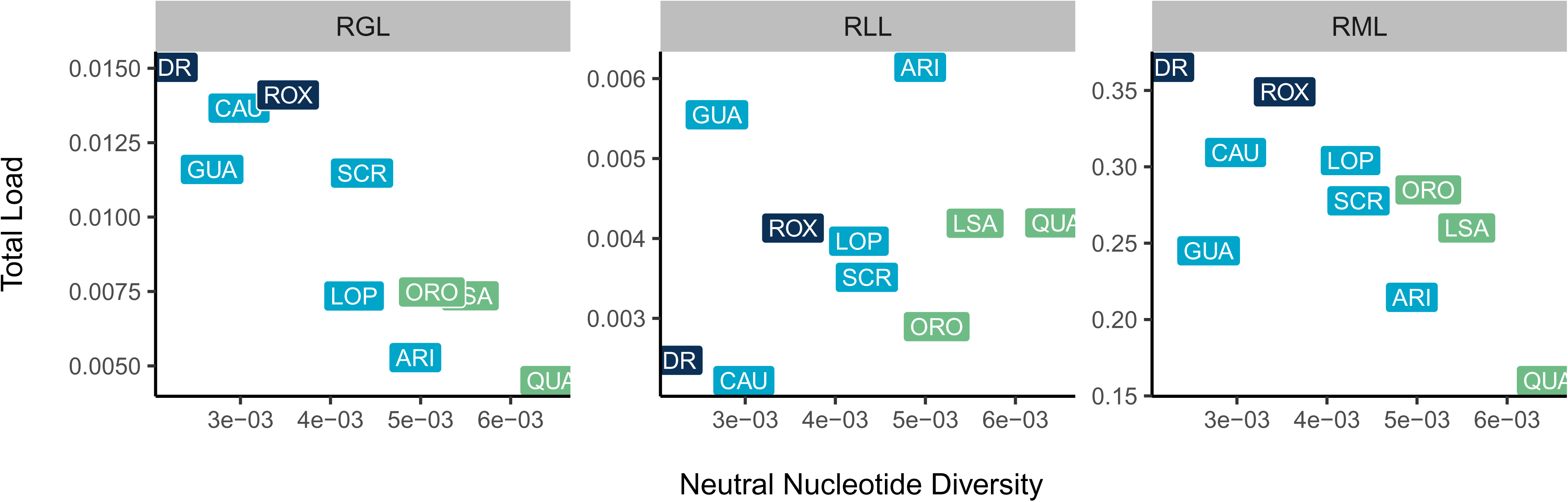
The relationship between neutral nucleotide diversity (calculated in four-fold degenerated sites) and per-population estimates of relative genetic load averaged within populations.

For the two rivers with samples suitable for analysis, Oropouche and Quare, our comparisons of upstream versus downstream populations agreed with previous studies (Willing et al. 2010; Qiu et al. 2022), and generally support lower effective sizes and/or bottlenecks experienced during upstream colonization. As expected, guppies from upstream sites tended to have higher load by almost all measures, except for heterozygous RLL (Supplemental Table S7 and Supplemental Fig. S3).

### Genetic load in the translocated and expanding population

To test hypothesis H1, we compared Guanapo (source) and upper Turure (translocated) populations. The results showed a complex pattern of differences in genetic load (Supplemental Table S8, Fig. 4). The upper Turure had significantly lower values of total and heterozygous RGL and RML, but higher homozygous RML than the Guanapo. For RLL, however, the upper Turure had significantly lower load on all three measures than the Guanapo.

**Figure 4.**
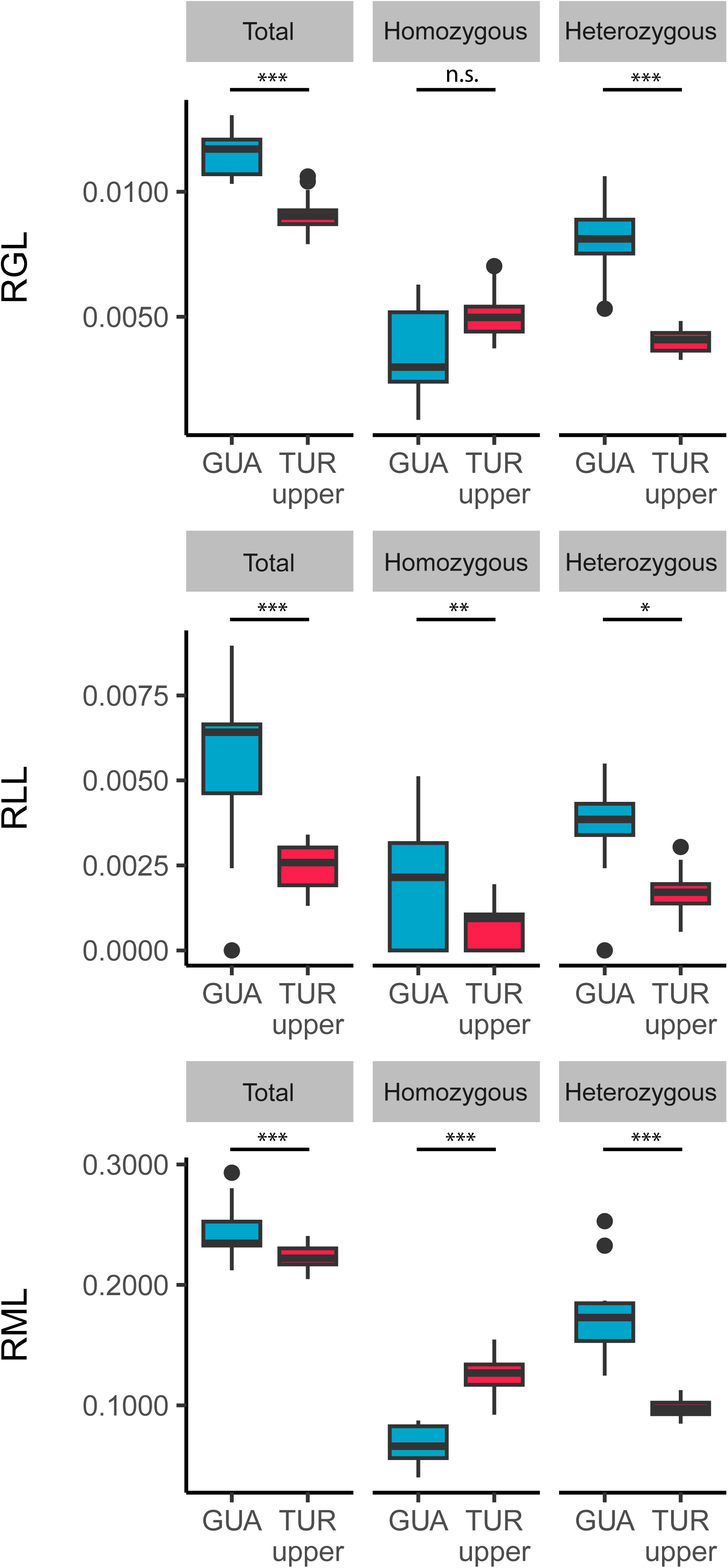
Comparisons of load estimates between Guanapo and upper Turure – pre- and post-translocation populations respectively. The top panel represents relative genetic load (RGL), the middle panel relative loss-of-function load (RLL) and the bottom panel relative missense load (RML). Boxplots are drawn from individual-based estimates with the horizontal line representing the population median, the boxes the interquartile range and the whiskers the maximum and minimum values. Horizontal lines with asterisks indicate statistically significant differences.

Focusing on the three Turure populations (hypothesis H2), we found that upper Turure had significantly higher RGL and RML compared to downstream populations, except for heterozygous RML and heterozygous RGL, which did not differ significantly from those of lower Turure guppies (Supplemental Table S9, Fig. 5). Remarkably, the pattern was reversed for RLL: upper Turure had significantly lower total, heterozygous and homozygous RLL compared to lower Turure, and also marginally lower homozygous RLL compared to middle Turure (Supplemental Table S9, Fig. 5).

**Figure 5.**
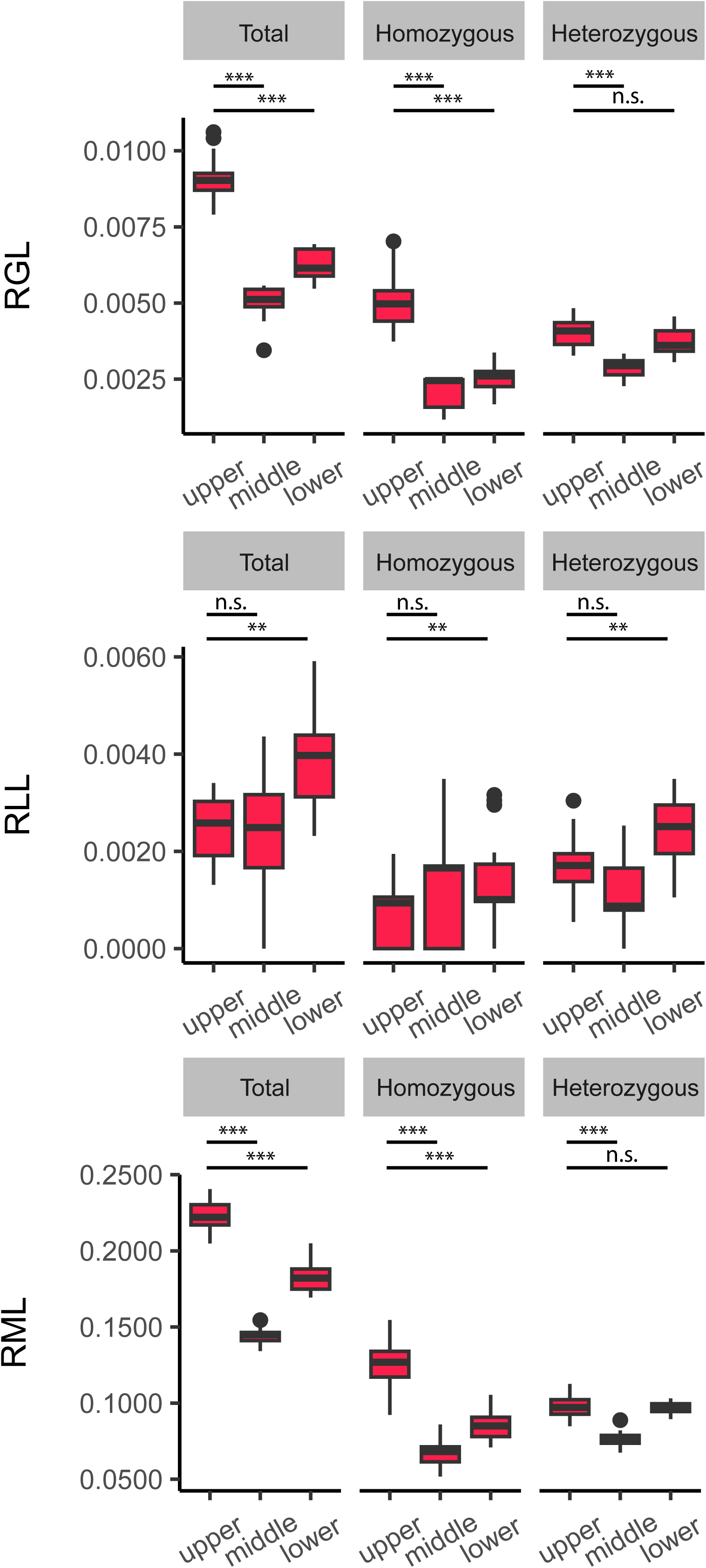
Comparisons of load estimates between the upper, middle and lower sites in the Turure river. The top panel shows relative genetic load (RGL), the middle panel shows relative loss-of-function load (RLL) and the bottom panel shows relative missense load (RML). Boxplots drawn from individual-based estimates with horizontal lines representing the population median, boxes the interquartile range, and whiskers the maximum and minimum values. Horizontal lines with asterisks indicate statistically significant differences.

## DISCUSSION

The accelerating pace of global change, driven by climate shifts and habitat loss, has led to widespread species migration, range fragmentation, and population isolation (Johnson et al. 2017). These rapid demographic shifts can profoundly influence the dynamics of genetic load, posing challenges for conservation biology. To date, most research in this area has centered on endangered, highly vulnerable, or even extinct taxa (Kuang et al. 2020; Von Seth et al. 2021; Kleinman-Ruiz et al. 2022; Ochoa et al. 2022; Xie et al. 2022). The role of genetic load in biological invasions has received less attention (de Pedro et al. 2021; Rougemont et al. 2020; Tayeh et al. 2013). Our study addresses this gap by investigating how genetic load accumulates in invasive, non-threatened species (Deacon et al. 2011; Lindholm et al. 2005; Rosenthal et al. 2021), thereby offering insights into the mechanisms of invasion—particularly when accompanied by admixture with native populations. Our large population-genomic dataset of hundreds of guppy genomes from multiple wild populations shows that genetic loads are strongly affected by historical effective population size (inferred from neutral diversity). We also found that translocated populations appear to have purged strongly deleterious mutations, and that subsequent admixture along the expansion gradient restored this load.

### Genetic load correlates with historical effective population sizes

Population genetics and conservation biology have long focused on how changes in population sizes shape the amount and distribution of deleterious variation (Bertorelle et al. 2022; van Oosterhout et al. 2026; Dussex et al. 2023; Robinson et al. 2023; Kimura et al. 1963; Lande 1998; Lynch et al. 1995). Theoretical and empirical evidence suggest that prolonged reductions in effective population size, or bottlenecks lasting hundreds or thousands of generations, should allow weakly to moderately deleterious alleles to accumulate, although strongly deleterious alleles might also be purged (Xue et al. 2015; Robinson et al. 2018; Dussex et al. 2021; Mathur and DeWoody 2021; Kleinman-Ruiz et al. 2022). Therefore, when measured based on mildly deleterious alleles, genetic load estimates should correlate with long-term Ne (or neutral nucleotide diversity). Testing these predictions using inter-species comparisons has produced mixed results: although some studies reported the expected correlations (Schmidt et al. 2023; Wilder et al. 2023), another reported no correlation (Wooldridge et al. 2024). However, no consistent general framework exists for comparing genetic load across species (Dussex et al. 2023; Teixeira and Huber 2021). Further, endangered species - the typical focus of such work - rarely contain multiple populations with contrasting demographic histories and a broad range of effective population sizes. Intraspecific associations between Ne and genetic load have thus been studied in only a handful of cases (Kleinman-Ruiz et al. 2022; Mathur and DeWoody 2021; Smeds et al. 2024; Mathur et al. 2023a; Grossen et al. 2020). Our intraspecific analyses therefore provide valuable insight, showing that the expected negative relationship between neutral nucleotide diversity and relative genetic load is detectable—particularly when comparing a broad spectrum of mutation types, as estimated by RGL and RML. Additionally, a significant relationship increased our confidence in the methods used to infer genetic load. These methods rely on predicted mutation effects and thus depend on the quality of annotation. Annotation used here is based on an ensemble pipeline and should be free of major issues, however, even in model species loss-of-function mutations tend to be enriched for annotation errors (Karczewski et al. 2020). While more sophisticated approaches for assessing variant impact are available, (Avsec et al. 2025; Cheng et al. 2023; Speak et al. 2023), they require large training datasets and are still rarely applied in conservation genomics. Our results demonstrate that methods based on simpler conservation scores can provide useful tools for wildlife population genomics.

### Purging in translocated populations of invasive potential

Our conclusion that the post-bottleneck Turure upper population experienced purging is based on distinct patterns observed for RLL, the load reflecting the most strongly deleterious mutations, in comparison with load estimates based on wider ranges of mutation effects (RML and RGL). A population bottleneck alone does not affect expected frequencies of variants, but increases variance in these frequencies and thus increases homozygosity. Consequently, under increased genetic drift alone, homozygous load is predicted to increase while heterozygous load should decline, and total genetic load should remain unchanged assuming intermediate effects of all variants in heterozygotes (Dussex et al. 2023; García-Dorado 2012). We detected the expected changes in heterozygous and homozygous loads when comparing source and translocated populations’ RML and RGL values (Fig. 4), though the total values have decreased in the translocated populations, possibly due to selection purging the most deleterious variants in homozygous genotypes. As expected (Glémin 2003; Caballero et al. 2017; Dussex et al. 2023), the bottleneck effect is most pronounced for strongly deleterious mutations: all three classes of RLL (total, heterozygous and homozygous) decreased following translocation. We acknowledge that the comparison of the Guanapo source population and the Turure translocated populations may be confounded by batch effects, as we used publicly available data from Guanapo. However, we controlled for sequencing depth in our analyses, and the batch effect that could have resulted from other sources was generally weak in the remaining populations. Furthermore, any bias would likely affect all classes of mutations similarly, synonymous ones including; therefore, given that our genetic load estimates are relative to synonymous ones, we do not expect this to substantially affect our overall results or interpretation. Finally, observation of different patterns using different classes of inferred mutations, as discussed above, cannot be reconciled with batch effect.

Effects on loads after bottlenecks similar to those we found have been documented in other species (Grossen et al. 2020; Khan et al. 2021; Quinn et al. 2024; Ochoa and Gibbs 2021; Xue et al. 2015; Stuart et al. 2025). Collectively, these findings provide strong evidence for purging (Dussex et al. 2023), suggesting that, under suitable conditions, highly deleterious mutations can be removed from bottlenecked populations, potentially increasing their invasiveness (Estoup et al. 2016).

The Turure population, which we infer to be purged of the most strongly deleterious mutations, exhibits marked invasive potential, a possibility first raised when indigenous guppies were observed to be displaced (Becher and Magurran 2000; Shaw et al. 1992). Early genetic studies using allozymes confirmed that the translocated individuals established a viable population, overcame the natural waterfall barrier—likely aided by flooding events—and began spreading into different downstream environments (Shaw et al. 1992). Subsequent analyses of a panel of SNPs showed that fish from the middle reaches of the Turure clustered with the Caroni drainage (Willing et al. 2010), and more recent marker-based studies distinguishing local from invasive individuals demonstrated that the invasion extended to, or beyond, the Turure–Quare confluence (Sievers et al. 2012). Continuous migration from above the barrier due to stochastic events such as flooding, together with subsequent effective mixing might explain this pattern (Sievers et al. 2012). However, the rapidity of the replacement of native fish in the downstream population, despite their larger effective population sizes, suggests that additional factors might have contributed to the success of the invaders. If so, the translocation experiment—designed to test the adaptive potential of guppies in novel environments—may ultimately have facilitated the loss of multiple populations within the drainage that were genetically distinct from guppies in other systems (Sievers et al. 2012).

The absence of strong geographic barriers in the lower Oropouche drainage likely promotes gene flow, which is particularly evident in the Quare population and, to a lesser extent, in La Seiva (Fig. 1A). However, our analyses revealed no evidence of Turure ancestry in the lower Quare or La Seiva populations (Fig. 1A), consistent with findings ∼15 years earlier (Willing et al. 2010). Nonetheless, the Turure invasion proceeded rapidly, and our results suggest that the initial bottleneck—by increasing the efficiency of purging strongly deleterious mutations—may have contributed to this successful colonization. Other studies of bottlenecked vertebrate populations have yielded mixed conclusions ranging from no evidence of purging at all (Robinson et al. 2019; Ochoa et al. 2022; Smeds et al. 2024), purging of only the most deleterious class of mutations (Grossen et al. 2020; Khan et al. 2021; Quinn et al. 2024), or accumulation of all classes of deleterious mutations (van der Valk et al. 2019). Disentangling whether such variation reflects methodological differences, limited statistical power, annotation quality, or true evolutionary processes remains difficult (Quinn et al. 2024). One critical factor may be the strength and duration of the bottleneck. Theory predicts that purging is most efficient under moderate reductions in population size, which expose recessive deleterious alleles to selection yet still allow selection to overcome genetic drift (Crow 1970; Glémin 2003; Robinson et al. 2023; Lombaert et al. 2025). Consistent with this, Quinn et al. (2024), found evidence of purging of loss-of-function alleles in populations that experienced less severe size declines, whereas populations undergoing the most rapid and drastic bottlenecks showed weaker signals of purging. This pattern suggests that extinction risk is the highest in severely reduced populations, where purging is the least effective. Our results align with these expectations. In the Turure River, the translocated population does not show a marked reduction in estimated neutral diversity or increase in Tajima’s D values, compared with its progenitor (Fig. 2, Supplemental Fig. S1). This likely reflects the moderate severity of the bottleneck and the relatively large number of individuals introduced (see Introduction; (Shaw et al. 1992; Sievers et al. 2012)). Despite these subtle changes, we nevertheless detect pronounced shifts in the genetic load estimates, supporting the view that purging can be effective under moderate bottlenecks.

### Admixture reshapes the landscape of genetic load in an expanding population

Our investigation of the Turure expansion gradient revealed that total RGL and RML decline further downstream (Fig. 5), which could, in principle, reflect repeated purging events following successive bottlenecks. However, this would cause a gradual loss of nucleotide diversity and a depletion of rare variants, while purging should primarily reduce heterozygous load by targeting the most deleterious mutations. In contrast, we found increased nucleotide diversity and lower Tajima’s D values in the middle and lower Turure populations compared with the up-river population. Moreover, the decline in genetic load is driven mainly by reductions in homozygous load, whereas heterozygous load remains largely unchanged or only moderately affected. Strikingly, for the most deleterious loss-of-function alleles, genetic load actually increases rather than decreases.

We therefore suggest that changes in genetic load along the Turure River primarily reflect admixture occurring below the waterfall barrier in the middle and lower reaches of the river. The upper and lower populations indeed differ substantially in their genetic composition, with the lower populations showing pronounced admixture with other ancestries (Fig. 1A). Depending on the genetic makeup of the admixed sources and the degree of divergence between them, such admixture could have profoundly reshaped the landscape of genetic load by increasing the heterozygote frequencies and introducing new haplotypes into the population.

Our analyses revealed a strong correlation between total RGL and long-term effective population size, as inferred from neutral nucleotide diversity (Fig. 3). Rivers in the Oropouche drainage generally exhibit higher nucleotide diversity than those in the Caroni drainage (Fig. 2), suggesting that a lower total RGL should be expected in Oropouche populations. This pattern is also supported in our comparisons of total RGL between the two drainages (Supplemental Table S5). If the native Turure ancestry from the Oropouche drainage followed this trend, it would have carried fewer moderately deleterious mutations than the Caroni guppies introduced during translocation. Consequently, admixture between these ancestries should, on average, reduce the genetic load of the expanding population consistent with our observations. As expected, this reduction in total RGL (and similarly in total RML) is driven primarily by decreases in homozygous load, likely reflecting the masking of moderately deleterious alleles that became fixed in the Caroni lineage after the Caroni-Oropouche divergence.

The most deleterious LoF mutations should rarely be fixed in unthreatened natural populations. Our neutral diversity dataset did not predict changes in RLL, suggesting that strongly deleterious mutations segregate and fix at comparable rates in different guppy populations (except for the purging in the Turure upriver population explained above). We also detected no significant difference in RLL between the Oropouche and Caroni drainages (Supplemental Tab. S5). Admixture is therefore the most likely explanation for an increase in RLL in the lower compared to upper Turure, which likely counteracted purging by increasing heterozygosity, at the same time introducing new deleterious mutations from an unpurged native Turure population (Fig. 5).

Overall, our results indicate that the recently translocated guppy population purged strongly deleterious mutations during the bottleneck, while the genetic load has been largely reestablished by subsequent mixing with local populations along the expansion gradient. Invader populations can thus exhibit a reshaped landscape of genetic load, with both population size reductions and admixture exerting significant influence on these patterns. These findings are relevant to understanding genetic processes in invasive species, which in a globally connected world are recognized as one of the major threats to biodiversity (Faulkner et al. 2024; Wallingford et al. 2020). Although biologists have long proposed that evolutionary processes play a key role in invasion success (Estoup et al. 2016), it is only recently that the abundance of deleterious alleles has been considered. Our results highlight that while deleterious alleles accumulate over thousands of generations (Wootton et al. 2023; van der Valk et al. 2019), they can be rapidly redistributed through genetic drift, selection, and gene flow when populations mix. Yet, it should be stressed that our analyses did not take into account several processes which may affect our interpretation, such as epistatic interactions, and local adaptation resulting from different environmental pressures encountered by the different populations we sampled. We emphasize that our study is correlational, and future studies should attempt to investigate the fitness consequences of the inferred genetic load. Furthermore, the temporal scope of our study reflects a relatively recent population expansion, and the long-term consequences of such demographic processes may differ over extended evolutionary timescales. Despite these limitations, our results highlight that admixture and biological invasions are associated with important consequences for genetic load segregating in populations.

## METHODS

### Sampling and sequencing

Individuals used in the study come from two batches of sampling and sequencing supplemented with 10 samples (Guanapo lower population) downloaded from the European Nucleotide Archive BioProject PRJEB10680. The first batch (51 individuals) of sampling was conducted in 2018 and sequenced in 2019. The second batch (140 individuals) was both sampled and sequenced in 2022. All individuals are wild fish from natural populations in Trinidad and Tobago. The sampling procedures complied with Polish law, European Directive 2010/63/EU and were conducted under permits issued by Ministry of Agriculture, Land and Fisheries of Trinidad and Tobago.

We sampled eight rivers on Trinidad - Aripo, Caura, Lopinot, and Santa Cruz (Caroni drainage), as well as La Seiva, Oropouche, Quare and Turure (Oropouche drainage) - and two rivers on Tobago: Dog River and Roxborough, collecting 191 individuals from 14 sampling locations/populations in total. Three rivers were sampled at multiple sites: Quare (upper and lower), Oropouche (upper and lower), and Turure (upper, middle and lower); see Fig 1B and Supplemental Table S1 for more details and geographic coordinates. Fish were anesthetized using tricaine mesylate (MS-222) and preserved in 98% ethanol. Tail fin tissue was used for DNA extraction (with Thermo Scientific™ MagJET™ Genomic DNA kit) and genomic libraries generation (with the NEBNext Ultra™ II FS DNA Library Prep Kit and NEBNext Multiplex Oligos for Illumina indices) according to manufacturer protocols. All samples were sequenced on the Illumina NovaSeq 6000 platform in 2x150bp mode at the NGI Sweden. Together with the ENA archive Guanapo samples, 201 samples from 15 populations were included for subsequent analyses.

### Population structure and nucleotide diversity

Raw sequencing reads were filtered with Trimmomatic (version 0.39; (Bolger et al. 2014)) and aligned to the guppy female reference genome (GCF_000633615.2; (Kunstner et al. 2016)) using bwa mem and default parameters (v0.7.10; (Li 2013)). The resulting BAM files were then subjected to duplicate marking with Picard tools (v2.21.6; http://broadinstitute.github.io/picard/). One sample from lower Oropouche had low mean coverage and was excluded from further analyses leaving, 200 samples in total.

To characterize the genetic composition of the samples, we performed principal component analysis (PCA) and ancestry inference. First, we jointly called variants in all BAM files using samtools mpileup (with -a AD parameter; v1.20, (Danecek et al. 2021)) and bcftools call ( with -G populations indicating parameter and -m multiallelic parameter; v1.20) Resulting VCF was then filtered with bcftools (version 0.1.12b, (Danecek et al. 2011)) using the following thresholds: MapppingQuality > 40, INFO/Depth < 5200, INFO/Depth > 1300, QUAL ≥ 30, MappingQualityBias > -3, ReadPositionBias < 3, SoftClipLengthBias < 3; and with vcftools (v0.1.12b, Danecek et al. 2011): MinorAlleleFraction ≥ 0.01, MaxAlelles = 2, MaxMissing = 50%; Depth ≥ 6 and GenotypeQuality ≥ 70. We then used plink (v1.9; (Purcell et al. 2007)) to transform the VCF file into a BED file and to prune it for variants in linkage disequilibrium using 200bp window, 1bp step and 0.5 r^2^ threshold. The resulting BED file was used to perform PCA. Using the same BED file, we ran Admixture (with the -s random seed parameter; v1.3, (Alexander et al. 2009)) 10 times for each K parameter (1-15). The Admixture results were plotted with Pong software (with -s 0.95 threshold to combine similar clusters at given K; v1.5, (Behr et al. 2016)).

To calculate neutral nucleotide diversity in our populations, we first identified fourfold-degenerated sites in the guppy genome using the find_4fds script (GitHub: https://github.com/mattheatley/extract_4fds). We then called genotypes in our samples using the same approach as above, including both polymorphic and monomorphic sites. The resulting VCF file was filtered with bcftools (v0.1.12b) as follows: MapppingQuality > 40, INFO/Depth < 5200, INFO/Depth > 1300, QUAL ≥ 30 and with vcftools (v0.1.12b): MaxAlelles = 2, MaxMissing = 50%; Depth ≥ 6 and retained only those that were located in 4-fold degenerated sites. Resulting files were used to calculate nucleotide diversity (π) in 1Mb windows across the genome with the Pixy tool (v1.2.7.beta1; (Korunes and Samuk 2021)). To estimate uncertainty around nucleotide diversity, we implemented a custom Python script that reproduced the estimator used in Pixy. For each population, individuals were resampled with replacement from the VCF genotype matrix, and nucleotide diversity was recalculated for each bootstrap replicate using the same pairwise comparison framework and missing-data handling as implemented in Pixy. This procedure was repeated 100 times to generate a bootstrap distribution of π estimates, from which the variance around the genome-wide estimate was derived.

### Ancestral allele inference and Conservation Scores

To infer guppy ancestral alleles (AA) we followed the approach of Smeds et al. (2024) and used genomic information from two closely related outgroups: *Micropoecilia picta* and *Xiphophorus maculatus*. Sequencing data for these two species were retrieved from NCBI’s SRA (accession numbers: ERR4077394 and SRR13649980, respectively) in FASTQ format. Reads were mapped to the guppy female reference genome and genotype calling was performed as described above, producing whole-genome VCF files These files were then used in the pseudo_haploidize.py script by Smeds et al. (2023, GitHub: https://github.com/linneas/wolf-deleterious) to infer ancestral alleles based on allele depth weight. Minimum depth thresholds were set to half of the expected coverage of *X. maculatus* and *M. picta* (7 and 11 respectively), and the minimum genotype quality was set to 30. Haplotypes that did not meet these thresholds were coded as “N”. The two resulting BED files (one per species) were merged and only sites where the nucleotide was identical for both species (and not N) were retained as informative for ancestral allele inference.

To obtain conservation scores (CS), we downloaded the CS BED file from the Ensembl fish Compara database (release 111, 65 fish species alignment, CS calculated with GERP, (Herrero et al. 2016)). Briefly, GERP quantifies evolutionary constraint by estimating the number of rejected substitutions at each alignment position. This is achieved by comparing the observed substitution rate to that expected under a neutral phylogenetic model inferred from the multiple sequence alignment. Positive GERP scores indicate sites under purifying selection (fewer substitutions than expected), whereas negative scores indicate an excess of substitutions relative to neutrality. The file was intersected with the ancestral allele BED file. The resulting CS-AA BED was used for genotype filtering, as described below.

### Genetic Load estimation

To estimate genetic load, we developed a bioinformatic pipeline analyzing within-population genomic variation. Each population was analyzed separately, enabling parallelization and the detection of rare variants. First, we inferred genotypes with GATK HaplotypeCaller (with -ERC BP_RESOLUTION and --output-mode EMIT_ALL_CONFIDENT_SITES parameters, v4.1.4.1; (McKenna et al. 2010)). Single individual genomic VCF (GVFCs) files were merged into population GVCFs (GATK CombineGVCFs) and used in GATK GenotypeGVCFs to produce VCF files in all-sites mode (-all-sites). Sites with fewer than 70% of genotypes meeting quality thresholds (DP ≥ 6 and GQ ≥ 30) were removed. Subsequently, we filtered out multiallelic sites and retained only sites with both CS and AA as one of the two alleles. Information about CS and AA was added to the INFO field using a custom Python script. In case of sites, where AA was the alternative to the reference nucleotide, the two alleles were swapped, so that REF always represents the ancestral allele in the final VCF file.

Because our samples differed in expected coverage, we applied individual-depth-based filtering. Only genotypes meeting the criteria 2 × mean DP of the individual ≥ DP ≥ 6 and GQ ≥ 30 were retained; genotypes failing these thresholds were coded as missing. Sites with more than 30% missing genotypes were excluded. Finally we extracted SNPs and removed variants failing GATK-recommended quality filters: QualityByDepth ≥ 2, MappingQuality ≥ 40, FisherScore ≤ 60, StrandOddsRatio ≤ 3, MappingQualityRankSum ≥ −12.5, ReadPosRankSum ≥ −8.

Per-individual relative genetic load (RGL) was calculated following the approach of Dussex et al. (2021), using the following formula:

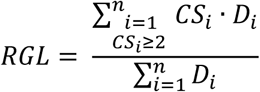

where n is the number of variants, CS_i_ is the conservation score and D_i_ is the number of derived alleles of the i^th^ variant. Only alleles at highly conserved sites (conservation score ≥ 2) were included in the numerator.

Additionally, we used called SNPs to identify loss of function (LOF, nonsense), missense and synonymous variants with SNPEff tool (v5.0; (Cingolani et al. 2012)) using built-in annotation originated from ensemble release 99. The values were calculated and normalized against synonymous derived alleles to calculate relative LOF load (RLL) using the following formulas:

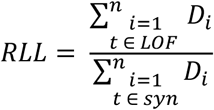

where n is the number of variants, t is the variant type and D_i_ is the number of derived alleles of the i^th^ variant.

Similarly, values for missense alleles were also used to calculate relative missense load (RML) using the following formula:

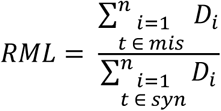

where n is the number of variants, t is the variant type and D_i_ is the number of derived alleles of the i^th^ variant.

The calculations were done separately for all variants (as a proxy for total genetic load), heterozygous genotypes (heterozygous genetic load) and homozygous genotypes (homozygous genetic load).

### Statistical analyses

All statistical analyses were performed in R (v4.5.3) Model assumptions were verified by examining diagnostic plots of residuals (or simulated residuals where appropriate) using the DHARMa package (Hartig 2016).

The association between mutation load and nucleotide diversity, was tested using general or generalized linear models, with a Gaussian distribution for RGL, and binomial distribution for RML and RLL. A logarithmic transformation was used to address the detected heteroskedasticity (reported in Supplemental Table S6). We initially fitted geographic region (Caroni, Oropuche, Tobago) in this model along with population, but the region was not significant in any analysis and some models containing it did not conform to model assumptions (heteroskedasticity), which improved when the region was removed. Therefore, we removed the region from all models.

We used general or generalized linear mixed models using glmmTMB package to compare genetic load between region, with population as a random effect and coverage as a covariate. We used a Gaussian distribution for RGL, and a binomial distribution for RML and RLL. We compared populations within Turure, between Turure and Guanapo and between the upper and lower parts of streams within Oropuche using general (RGL) or generalized (RGL and RLL) linear models. These models included coverage as a covariate and batch as a fixed effect (except for the Turure and Guanapo comparison, in which batch and population effects could not be differentiated). For the generalized models, we used a binomial distribution, or, when over- or underdispersion was detected, a quasibinomial distribution (in which case we report the dispersion parameter, Supplemental Tab. S7 – S9). In our analysis of the differences between regions (Supplemental Tab. S5) in nucleotide diversity (Supplemental Tab. S6), only the lower river sites were used and all Turure populations were excluded.

## Declaration of generative AI in the writing process

During the preparation of this manuscript we used ChatGPT to improve the language and clarity of the text. Authors reviewed and edited the content after using this tool, and take full responsibility for the final version of the manuscript.

## DATA ACCESS

All raw sequencing data generated in this study have been submitted to the NCBI Short Read Archive under accession number PRJNA1040274.

All codes are available in a GitHub repository (https://github.com/0-Ioniel-0/Expansion_Load) and in the Supplemental Code.

## ACKNOWLADGMENTS

We are grateful to Deborah Charlesworth, Wiesław Babik and Jonathan Parrett for their valuable comments on an early version of the manuscript, and to the members of the Evolutionary Biology Group for their insightful discussions and continuous support. We thank Katarzyna Dudek from Jagiellonian University for preparing genomic libraries. This work was funded by the Polish National Science Centre 2018/31/D/NZ8/00091 grant. M.K. was supported by the Polish National Agency for Academic Exchange under the Bekker Programme. Computations were performed at the Poznan Supercomputing and Networking Centre. We are also grateful to the Fisheries Division of the Ministry of Agriculture and Fisheries, Republic of Trinidad and Tobago for granting permission to collect and export of fish tissue.

