## Supplemental Material for "Interplay between purging and admixture shapes genetic load in an invasive guppy population"

### Supplementary Figures

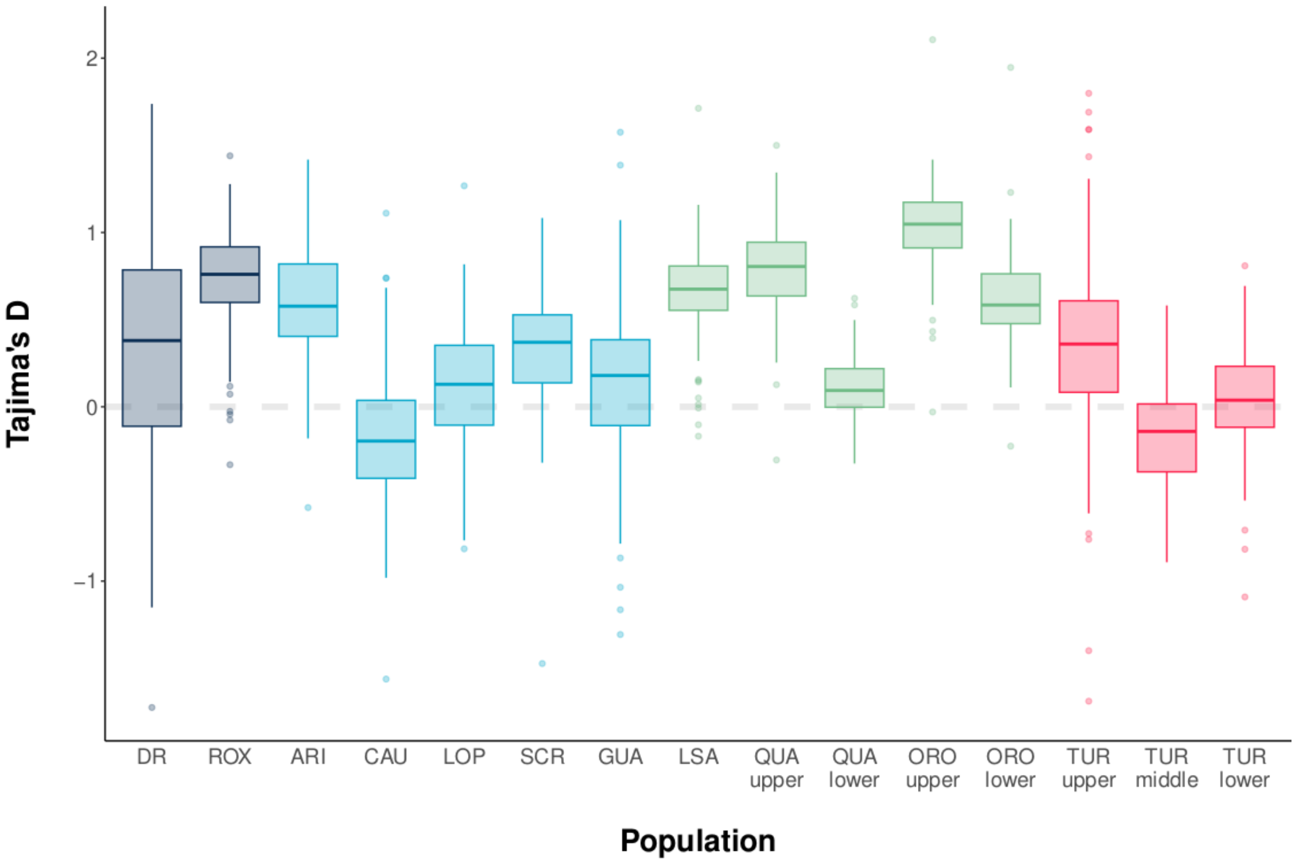


**Fig. S1.** Tajima's D in studied populations. The regions are color-coded as follows: Tobago (dark blue), Caroni drainage (light blue), Oropouche drainage (green), Turure river (red).

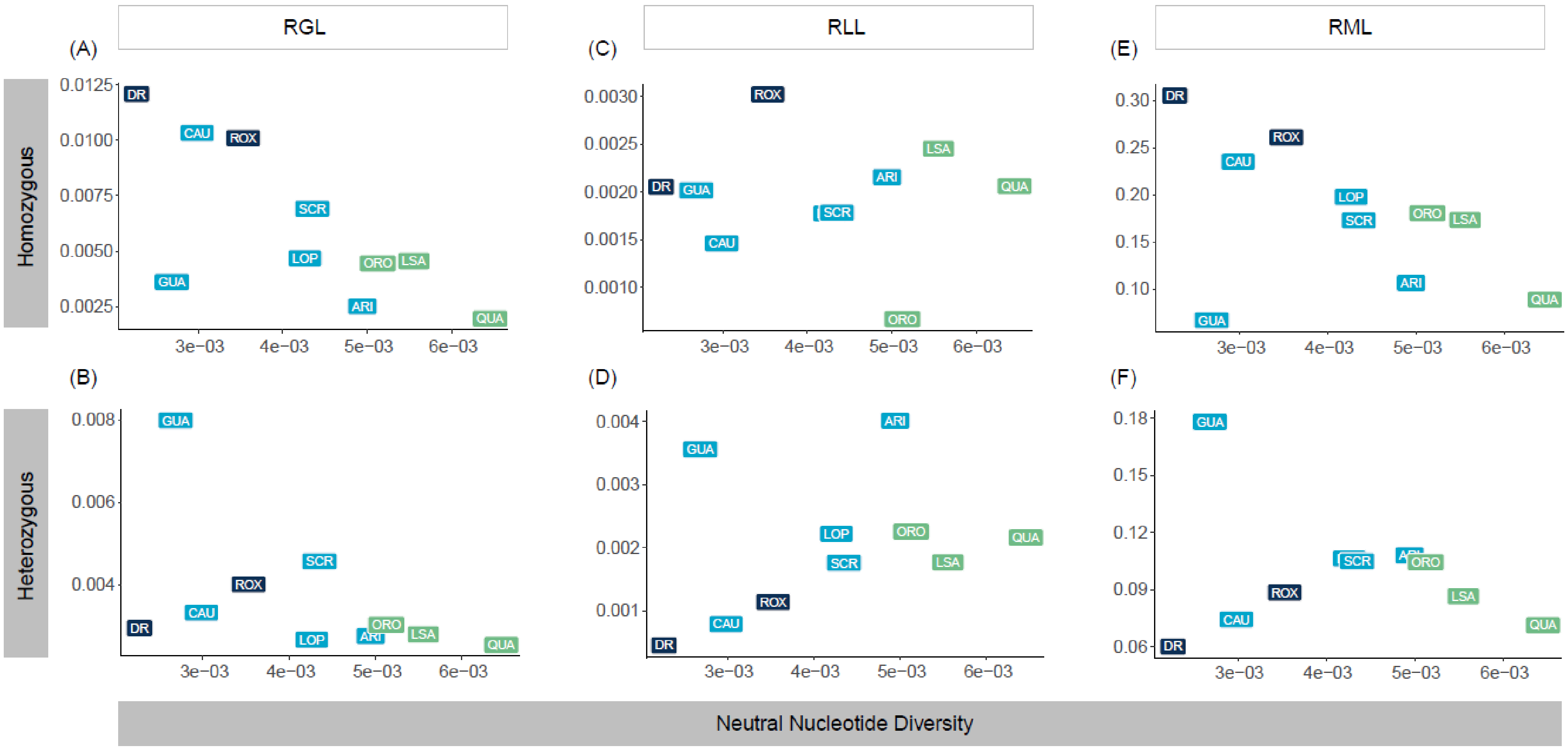


**Fig. S2.** Relation between neutral nucleotide diversity and three measures of genetic load in homozygote and heterozygote genotypes. Left panel represents relative genetic load (RGL), middle panel relative loss-of-function load (RLL) and right panel relative missense load (RML). The regions are colour-coded as follows: Tobago (dark blue), Caroni (light blue) and Oropuche (green).

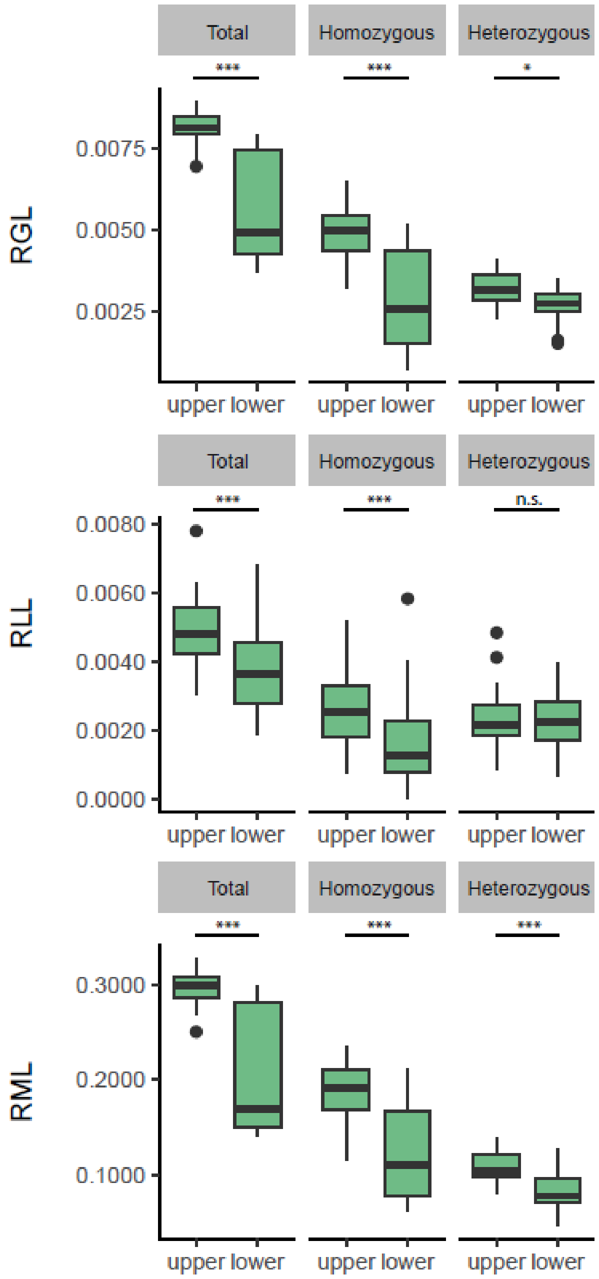


**Fig. S3.** Comparisons of load estimates between upper and lower sites of Oropouche and Quare rivers. Top panel represents relative genetic load (RGL), middle panel relative loss-of-function load (RLL) and bottom panel relative missense load (RML Boxplots drawn from individual-based estimates with horizontal line representing population median, boxplot interquartile and whiskers span maximum and minimum values. Horizontal lines with asterixis indicate statistically significant differences.

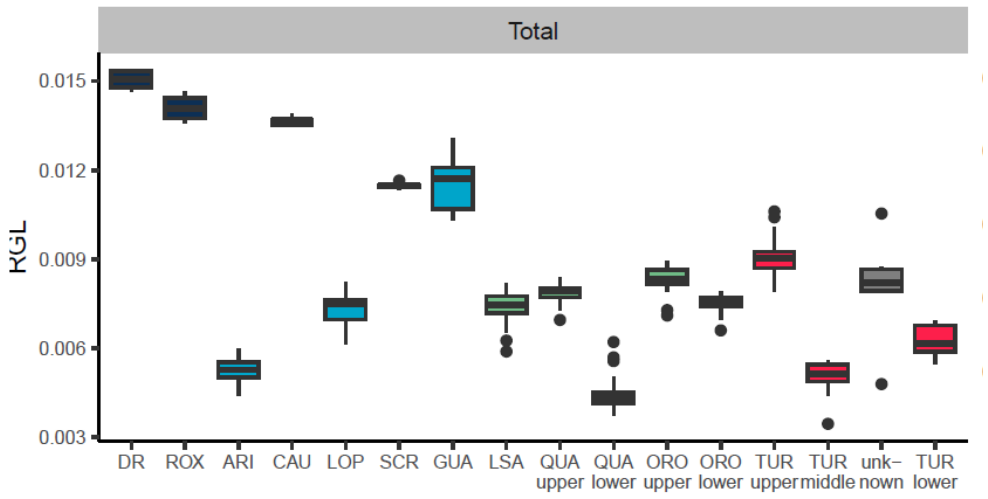


**Fig. S5.** Total relative genetic load in all populations. The regions are color-coded as follows: Tobago (dark blue), Caroni (light blue), Oropouche (green), and Turure (red). Gray color shows RGL in individuals of unknown ancestry sampled in Turure middle site.

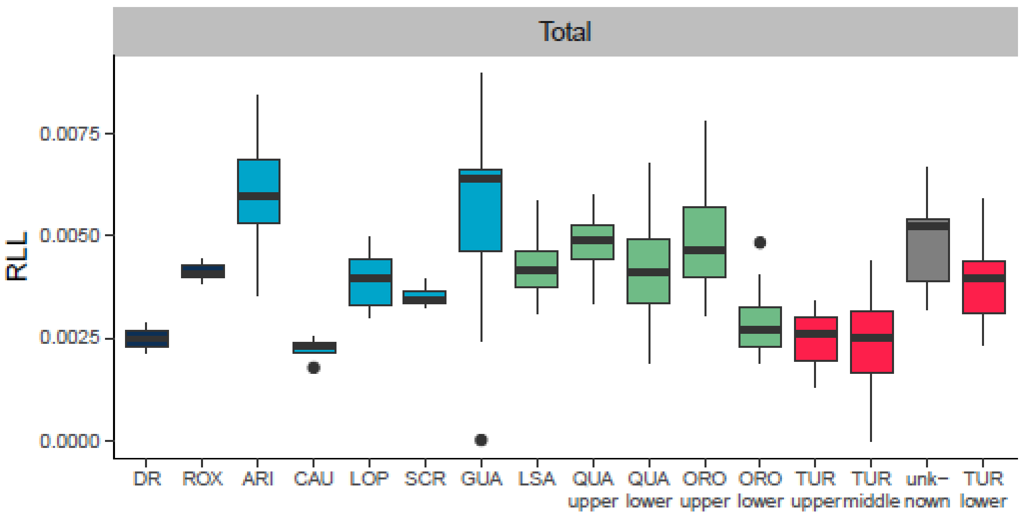


**Fig. S6.** Total loss-of-function genetic load in all populations. The regions are color-coded as follows: Tobago (dark blue), Caroni (light blue), Oropouche (green), and Turure (red). Gray color shows RGL in individuals of unknown ancestry sampled in Turure middle site.

### Supplementary Tables

**Table S1** Sampling sites on Trinidad and Tobago. The first column is site abbreviation, the 5th, 6th and 7th column is total number of individuals used in this study.

| Population | Island | Drainage | Location | N | Males | Females | Juveniles |
|---|---|---|---|---|---|---|---|
| DR | Tobago | East flowing | N 11.22556 W 060.64500 | 5 | 1 | 4 | - |
| ROX | Tobago | East flowing | N 11.25056 W 060.58250 | 5 | 2 | 3 | - |
| ARI | Trinidad | Caroni | N 10.66969 W 061.22991 | 14 | 8 | 6 | - |
| CAU | Trinidad | Caroni | N 10.70797 W 061.35759 | 5 | 2 | 3 | - |
| LOP | Trinidad | Caroni | N 10.69333 W 061.32195 | 19 | 7 | 12 | - |
| SCR | Trinidad | Caroni | N 10.71565 W 061.46842 | 5 | 2 | 1 | 2 |
| LSA | Trinidad | Oropouche | N 10.65718 W 061.15276 | 19 | 8 | 11 | - |
| QUA upper | Trinidad | Oropouche | N 10.67610 W 061.19606 | 11 | 1 | 6 | 4 |
| QUA lower | Trinidad | Oropouche | N 10.65265 W 061.18951 | 22 | 7 | 12 | 3 |
| ORO upper | Trinidad | Oropouche | N 10.67048 W 061.13784 | 20 | 3 | 3 | 14 |
| ORO lower | Trinidad | Oropouche | N 10.65963 W 061.13137 | 13 | 6 | 7 | - |
| TUR upper | Trinidad | Oropouche | N 10.67747 W 061.16355 | 19 | 8 | 11 | - |
| TUR middle | Trinidad | Oropouche | N 10.65687 W 061.16776 | 19 | 4 | 15 | - |
| TUR lower | Trinidad | Oropouche | N 10.61899 W 061.15423 | 14 | 7 | 4 | 3 |

**Tab. S2.** Number of sites remained after each filtering step of the pipeline.

| Population | Mapped Scaffolds | After Pre-Filtering (PF) | PF Removed | PF Removed Fraction | After AACS | AACS Removed | AACS Removed Fraction | After Final-Filtering (FF) | FF Removed | FF Removed Fraction |
|---|---|---|---|---|---|---|---|---|---|---|
| ARI | 696 700 953 | 510 842 293 | 185 858 660 | 26.68% | 43732590 | 467109703 | 91.44% | 23 631 720 | 20 100 870 | 45.96% |
| CAU | 696 700 953 | 420 109 925 | 276 591 028 | 39.70% | 47 085 196 | 373 024 729 | 88.79% | 23 425 829 | 23 659 367 | 50.25% |
| DR | 696 700 953 | 403 232 383 | 293 468 570 | 42.12% | 44 697 914 | 358 534 469 | 88.92% | 22 202 604 | 22 495 310 | 50.33% |
| GUA | 696 700 953 | 621 006 288 | 75 694 665 | 10.86% | 12 343 749 | 608 662 539 | 98.01% | 226 410 | 12 117 339 | 98.17% |
| LOP | 696 700 953 | 437 206 382 | 259 494 571 | 37.25% | 42 441 084 | 394 765 298 | 90.29% | 12 538 454 | 29 902 630 | 70.46% |
| LSA | 696 700 953 | 502 537 963 | 194 162 990 | 27.87% | 50 665 986 | 451 871 977 | 89.92% | 25 675 105 | 24 990 881 | 49.32% |
| ORO lower | 696 700 953 | 317 185 665 | 379 515 288 | 54.47% | 29 923 754 | 287 261 911 | 90.57% | 13 964 074 | 15 959 680 | 53.33% |
| ORO upper | 696 700 953 | 375 051 056 | 321 649 897 | 46.17% | 33 266 992 | 341 784 064 | 91.13% | 17 946 984 | 15 320 008 | 46.05% |
| QUA lower | 696 700 953 | 308 417 031 | 388 283 922 | 55.73% | 26 606 346 | 281 810 685 | 91.37% | 14 602 381 | 12 003 965 | 45.12% |
| QUA upper | 696 700 953 | 271 245 660 | 425 455 293 | 61.07% | 23 166 424 | 248 079 236 | 91.46% | 11 717 456 | 11 448 968 | 49.42% |
| ROX | 696 700 953 | 586 243 273 | 110 457 680 | 15.85% | 65 926 450 | 520 316 823 | 88.75% | 32 902 050 | 33 024 400 | 50.09% |
| SCR | 696 700 953 | 376 645 690 | 320 055 263 | 45.94% | 43 910 262 | 332 735 428 | 88.34% | 21 828 512 | 22 081 750 | 50.29% |
| TUR lower | 696 700 953 | 401 211 576 | 295 489 377 | 42.41% | 41 060 442 | 360 151 134 | 89.77% | 20 509 478 | 20 550 964 | 50.05% |
| TUR middle | 696 700 953 | 352 095 559 | 344 605 394 | 49.46% | 36 165 812 | 315 929 747 | 89.73% | 18 026 588 | 18 139 224 | 50.16% |
| TUR upper | 696 700 953 | 346 896 822 | 349 804 131 | 50.21% | 36 023 210 | 310 873 612 | 89.62% | 17 957 822 | 18 065 388 | 50.15% |

*Note: PF – pre-filtering based on 30% missing individuals allowed (DP <= 6 and GQ <= 30 also counting as missing, but not marked at this point); AACS – filtering based on presence of Ancestral Allele and Conservation Score; FF – final filtering based on 30% missing individuals allowed (2x mean DP <= DP <= 6 and GQ <30 also marked as missing - changed to ./.).*

**Table S3.** Number of analyzed variants across all populations by annotation and Conservation Scores (CS) categories.

| Category | Total | CS>0 | CS>1 | CS>2 | CS>3 | CS>4 |
|---|---|---|---|---|---|---|
| Intergenic | 318974 | 57346 | 3744 | 544 | 33 | 3 |
| Intronic | 794424 | 141706 | 7594 | 926 | 68 | 1 |
| UTR | 77233 | 14086 | 1145 | 154 | 11 | 0 |
| Missense | 13954 | 5618 | 1835 | 775 | 145 | 6 |
| Nonsense | 45 | 19 | 2 | 1 | 0 | 0 |
| Synonymous | 90851 | 9779 | 2938 | 580 | 34 | 0 |
| HIGH | 287 | 87 | 33 | 7 | 1 | 0 |
| MODERATE | 14242 | 5752 | 1885 | 797 | 149 | 6 |
| LOW | 107072 | 12336 | 3344 | 665 | 43 | 0 |
| MODIFIER | 1537782 | 275926 | 15026 | 2010 | 151 | 4 |
| Total | 1659383 | 294101 | 20288 | 3479 | 344 | 10 |

**Table S4.** Population genomic summary statistics and functional variant annotation**.**

| Population | Cov. | Nonref alleles | Het | Sing. | Ti/Tv | HOM CS>0 | HET CS>0 | HOM CS>1 | HET CS>1 | HOM CS>2 | HET CS>2 | HOM CS>3 | HET CS>3 | HOM CS>4 | HET CS>4 | HET HIGH | HOM HIGH | HET MOD. | HOM MOD. | HET LOW | HOM LOW |
|---|---|---|---|---|---|---|---|---|---|---|---|---|---|---|---|---|---|---|---|---|---|
| ARI | 15.1 | 45916.2 | 31747.5 | 223.1 | 1.48 | 9439 | 12490 | 454 | 695 | 77 | 101 | 7 | 9 | 2 | 2 | 7.1 | 1.9 | 192.5 | 95.4 | 1265.3 | 260.9 |
| CAU | 14.3 | 78809.4 | 30157.4 | 3084.0 | 1.63 | 17655 | 16494 | 1373 | 1307 | 271 | 240 | 33 | 23 | 1 | 0 | 5.4 | 5.0 | 509.0 | 811.2 | 2613.6 | 2133.6 |
| DR | 13.7 | 114272.2 | 24475.6 | 2041.0 | 1.64 | 27733 | 16124 | 2063 | 1317 | 414 | 273 | 51 | 30 | 0 | 0 | 4.2 | 10.2 | 589.4 | 1510.0 | 2135.8 | 3867.8 |
| GUA | 10.7 | 30623.8 | 16628.6 | 222.8 | 1.49 | 4983 | 9008 | 288 | 549 | 53 | 100 | 5 | 11 | 0 | 0 | 6.1 | 1.5 | 142.0 | 31.6 | 804.7 | 289.8 |
| LOP | 15.3 | 65692.7 | 32043.4 | 430.7 | 1.52 | 17374 | 20353 | 980 | 1197 | 154 | 177 | 9 | 12 | 0 | 0 | 6.4 | 2.6 | 309.6 | 294.9 | 1478.5 | 733.5 |
| LSA | 14.2 | 71435.2 | 38461.6 | 524.9 | 1.57 | 16732 | 19770 | 950 | 1203 | 163 | 201 | 21 | 23 | 1 | 1 | 7.2 | 4.9 | 351.2 | 355.8 | 2257.3 | 914.9 |
| ORO lower | 13.6 | 55176.8 | 27472.9 | 783.1 | 1.52 | 13884 | 15078 | 735 | 845 | 128 | 145 | 18 | 19 | 0 | 0 | 5.6 | 0.8 | 264.2 | 231.2 | 1375.3 | 585.6 |
| ORO upper | 13.5 | 56783.6 | 26790.2 | 203.9 | 1.53 | 16062 | 16412 | 910 | 966 | 153 | 163 | 22 | 20 | 0 | 0 | 4.7 | 3.4 | 235.8 | 247.5 | 1189.8 | 603.1 |
| QUA lower | 13.7 | 32614.0 | 23568.3 | 158.5 | 1.54 | 5650 | 8814 | 233 | 483 | 41 | 76 | 1 | 2 | 0 | 0 | 3.5 | 1.6 | 115.1 | 70.4 | 1185.9 | 216.8 |
| QUA upper | 12.8 | 55837.8 | 32880.2 | 973.4 | 1.52 | 13622 | 16348 | 692 | 884 | 135 | 161 | 15 | 16 | 1 | 1 | 7.4 | 2.6 | 321.1 | 207.9 | 1638.9 | 484.1 |
| ROX | 15.3 | 146280.0 | 48755.8 | 2339.6 | 1.65 | 33871 | 27611 | 2541 | 2194 | 509 | 443 | 59 | 54 | 2 | 2 | 13.6 | 18.4 | 1076.6 | 1589.8 | 4088.4 | 4051.0 |
| SCR | 13.6 | 83161.8 | 44891.6 | 2866.6 | 1.61 | 16611 | 19951 | 1180 | 1533 | 228 | 284 | 24 | 34 | 1 | 1 | 12.8 | 6.4 | 764.0 | 632.8 | 4031.0 | 1637.6 |
| TUR lower | 13.1 | 32661.9 | 25013.6 | 118.6 | 1.52 | 5545 | 8224 | 320 | 562 | 61 | 96 | 6 | 6 | 0 | 0 | 4.8 | 1.4 | 193.9 | 86.1 | 1550.9 | 223.6 |
| TUR upper | 13.2 | 27809.2 | 16955.3 | 72.6 | 1.55 | 6109 | 7686 | 367 | 550 | 69 | 104 | 10 | 12 | 1 | 1 | 3.6 | 0.8 | 207.3 | 134.8 | 1329.5 | 401.3 |
| TUR_middle | 13.4 | 20512.4 | 15154.5 | 66.5 | 1.52 | 3149 | 5062 | 160 | 347 | 34 | 56 | 2 | 2 | 1 | 1 | 1.0 | 1.1 | 83.8 | 49.1 | 789.2 | 141.7 |

*Note:* In the table, column represent different population statistics, as follow: **Cov.** - average sequencing depth across all genotypes in the population; **Nonref allels** - an average number of nonreference alleles per individual; **Het** - average number of heterozygous genotypes per individual; **Sing.** - an average number of singletons per individual; **Ti/Tv** - Transition/Transversion ratio; **CS columns** - number of loci where at least one individual is homozygous (HOM) or heterozygous (HET) with conservation scores (CS) larger than given number. **Impact statistics** - average number of heterozygous or homozygous variants per individual with given SnpEff impact statistics (HIGH, MODERATE, LOW).

**Table S5.** Comparison of genetic load (RGL: relative genetic load; RML: relative missense load; RLL: relative loss of function load) between regions, with estimate of Oropouche in the intercept. Reported statics are t for lms on RGL, z for glms for missense and loss of function. Significant P-values for the region effects are in bold.

| | | | | | | | Random effects | |
|---|---|---|---|---|---|---|---|---|
| **Dependent** | **Term** | **Estimate** | **SE** | **Statistic** | **P** | **Stat type** | **Variance** | **Std. Dev.** |
| Total RGL | Intercept | 0.0062 | 0.0014 | 4.59 | 0.000 | z | Population | |
| | Caroni | 0.0034 | 0.0017 | 2.02 | **0.043** | z | 5.4E-06 | 0.0023 |
| | Tobago | 0.0081 | 0.0021 | 3.83 | **<0.001** | z | | |
| | Coverage | 0.0000 | 0.0000 | 0.83 | 0.405 | z | | |
| ln(Heterozygous RGL) | Intercept | -5.7915 | 0.1775 | -32.63 | 0.000 | z | Population | |
| | Caroni | 0.3266 | 0.2098 | 1.56 | 0.120 | z | 8.1E-02 | 0.2839 |
| | Tobago | 0.2126 | 0.2645 | 0.80 | 0.422 | z | | |
| | Coverage | -0.0065 | 0.0042 | -1.56 | 0.118 | z | | |
| Homozygous RGL | Intercept | 0.0033 | 0.0013 | 2.58 | 0.010 | z | Population | |
| | Caroni | 0.0020 | 0.0015 | 1.28 | 0.200 | z | 4.3E-06 | 0.0021 |
| | Tobago | 0.0075 | 0.0019 | 3.87 | **<0.001** | z | | |
| | Coverage | 0.0000 | 0.0000 | 1.07 | 0.286 | z | | |
| Total RML | Intercept | -1.5148 | 0.1014 | -14.93 | 0.000 | z | Population | |
| | Caroni | 0.1647 | 0.1267 | 1.30 | 0.194 | z | 3.0E-02 | 0.1730 |
| | Tobago | 0.4543 | 0.1582 | 2.87 | **0.004** | z | | |
| | Coverage | 0.0022 | 0.0011 | 2.03 | **0.042** | z | | |
| Heterozygous RML | Intercept | -2.3653 | 0.1306 | -18.12 | 0.000 | z | Population | |
| | Caroni | 0.2264 | 0.1618 | 1.40 | 0.162 | z | 4.9E-02 | 0.2207 |
| | Tobago | -0.1729 | 0.2021 | -0.86 | 0.392 | z | | |
| | Coverage | -0.0058 | 0.0017 | -3.35 | **0.001** | z | | |
| Homozygous RML | Intercept | -2.0573 | 0.2106 | -9.77 | 0.000 | z | Population | |
| | Caroni | 0.0312 | 0.2654 | 0.12 | 0.906 | z | 1.3E-01 | 0.3630 |
| | Tobago | 0.7065 | 0.3315 | 2.13 | **0.033** | z | | |
| | Coverage | 0.0060 | 0.0012 | 4.81 | **<0.001** | z | | |
| Total RLL | Intercept | -5.6672 | 0.2040 | -27.78 | 0.000 | z | Population | |
| | Caroni | 0.0891 | 0.2082 | 0.43 | 0.669 | z | 7.3E-02 | 0.2702 |
| | Tobago | -0.1270 | 0.2571 | -0.49 | 0.621 | z | | |
| | Coverage | 0.0039 | 0.0080 | 0.49 | 0.625 | z | | |
| Heterozygous RLL | Intercept | -6.1015 | 0.3107 | -19.64 | 0.000 | z | Population | |
| | Caroni | 0.0424 | 0.3254 | 0.13 | 0.896 | z | 1.8E-01 | 0.4282 |
| | Tobago | -1.0564 | 0.4137 | -2.55 | **0.011** | z | | |
| | Coverage | -0.0069 | 0.0116 | -0.59 | 0.552 | z | | |
| Homozygous RLL | Intercept | -6.6876 | 0.2447 | -27.33 | 0.000 | z | Population | |
| | Caroni | 0.1427 | 0.2270 | 0.63 | 0.530 | z | 7.5E-02 | 0.2745 |
| | Tobago | 0.4796 | 0.2702 | 1.78 | **0.076** | z | | |
| | Coverage | 0.0150 | 0.0109 | 1.37 | 0.171 | z | | |

**Table S6.** Results of linear models testing the effect of population size, approximated with nucleotide diversity, on three measures of genetic load (RGL: relative genetic load; RML: relative missense load; RLL: relative loss of function load). Significant P-values for the effect of diversity are in bold.

| Dependent | Term | Estimate | SE | t | P |
|---|---|---|---|---|---|
| Total RGL | intercept | 0.020 | 0.002 | 9.778 | <0.001 |
| | diversity | -2.519 | 0.473 | -5.317 | **<0.001** |
| Heterozygous RGL | intercept | 0062 | 0.002 | 3.75 | 0.006 |
| | diversity | 0.605 | 0.376 | --1.609 | 0.146 |
| Homozygous RGL | intercept | 0.014 | 0.003 | 5.009 | 0.001 |
| | diversity | 1.914 | 0.642 | 2.977 | **0.018** |
| Total RML | intercept | 0.415 | 0.049 | 8.442 | <0.001 |
| | diversity | 32.753 | 11.163 | -2.934 | **0.019** |
| Heterozygous RML | intercept | 0.117 | 0.037 | 3.161 | 0.013 |
| | diversity | 4.532 | 8.423 | 0.538 | 0.605 |
| Homozygous RML | intercept | 0.297 | 0.076 | 3.90 | 0.004 |
| | diversity | 28.220 | 17.314 | --1.63 | 0.142 |
| Total RLL | intercept | 0.002 | 0.001 | 2.099 | 0.069 |
| | diversity | 0.238 | 0.316 | 0.753 | 0.473 |
| Heterozygous RLL | intercept | 0.001 | 0.001 | 0.690 | 0.510 |
| | diversity | 0.272 | 0.279 | 0.976 | 0.357 |
| Homozygous RLL | intercept | 0.002 | 0.001 | 2.927 | 0.019 |
| | diversity | 0.034 | 0.161 | -0.212 | 0.837 |

**Table S7.** Comparison of genetic load of lower (in the intercept) and upper sites in Oropouche (in the intercept), and Qare population, including coverage and sequencing batch as covariates. Reported statistics are t-values for linear models and z-values for generalized linear models. Significant P-values are shown in bold. When error distribution was over/under dispersed, quasibinomial distribution was fitted and dispersion parameter (d.p.) is reported in the first column.

| Dependent | Term | Estimate | SE | Statistic | P | Stat_type |
|---|---|---|---|---|---|---|
| Total RGL | Intercept | 0,0075 | 0,0006 | 12,79 | <0,001 | t |
| | Population | -0,0018 | 0,0002 | -9,05 | **<0,001** | t |
| | Site | 0,0019 | 0,0002 | 9,55 | **<0,001** | t |
| | Coverage | 0,0000 | 0,0000 | -1,13 | 0,264 | t |
| | Batch | -0,0006 | 0,0002 | -2,33 | **0,023** | t |
| Heterozygous RGL | Intercept | 0,0032 | 0,0004 | 8,67 | <0,001 | t |
| | Population | -0,0001 | 0,0001 | -0,60 | 0,552 | t |
| | Site | 0,0004 | 0,0001 | 3,27 | **0,002** | t |
| | Coverage | 0,0000 | 0,0000 | -1,13 | 0,262 | t |
| | Batch | -0,0001 | 0,0002 | -0,82 | 0,414 | t |
| Homozygous RGL | Intercept | 0,0043 | 0,0007 | 6,62 | <0,001 | t |
| | Population | -0,0017 | 0,0002 | -7,81 | **<0,001** | t |
| | Site | 0,0015 | 0,0002 | 6,75 | **<0,001** | t |
| | Coverage | 0,0000 | 0,0000 | -0,38 | 0,708 | t |
| | Batch | -0,0004 | 0,0003 | -1,64 | 0,107 | t |
| Total RML (d.p. 6,34 ) | Intercept | -1,1578 | 0,0962 | -12,04 | <0,001 | t |
| | Population | -0,2962 | 0,0303 | -9,77 | **<0,001** | t |
| | Site | 0,2548 | 0,0303 | 8,40 | **<0,001** | t |
| | Coverage | -0,0118 | 0,0059 | -2,02 | **0,048** | t |
| | Batch | -0,2070 | 0,0399 | -5,19 | **<0,001** | t |
| Heteroygous RML (d.p. 5,38) | Intercept | -2,1768 | 0,1354 | -16,07 | <0,001 | t |
| | Population | -0,0360 | 0,0416 | -0,87 | 0,390 | t |
| | Site | 0,1883 | 0,0426 | 4,43 | **<0,001** | t |
| | Coverage | -0,0136 | 0,0082 | -1,65 | 0,104 | t |
| | Batch | -0,1938 | 0,0560 | -3,46 | **0,001** | t |
| Homozygous RML (d.p. 7,49) | Intercept | -1,6033 | 0,1271 | -12,61 | <0,001 | t |
| | Population | -0,4590 | 0,0411 | -11,17 | **<0,001** | t |
| | Site | 0,2920 | 0,0402 | 7,26 | **<0,001** | t |
| | Coverage | -0,0112 | 0,0078 | -1,44 | 0,155 | t |
| | Batch | -0,2189 | 0,0532 | -4,12 | **<0,001** | t |
| Total RLL | Intercept | -6,0277 | 0,2037 | -29,60 | <0,001 | t |
| | Population | 0,1663 | 0,0634 | 2,62 | **0,011** | t |
| | Site | 0,3527 | 0,0665 | 5,30 | **<0,001** | t |
| | Coverage | 0,0187 | 0,0119 | 1,56 | 0,123 | t |
| | Batch | 0,0224 | 0,0830 | 0,27 | 0,788 | t |
| Heterozygous RLL | Intercept | -5,7490 | 0,2777 | -20,70 | <0,001 | t |
| | Population | 0,1865 | 0,0831 | 2,25 | **0,028** | t |
| | Site | 0,0240 | 0,0861 | 0,28 | 0,781 | t |
| | Coverage | -0,0280 | 0,0169 | -1,66 | 0,102 | t |
| | Batch | -0,2266 | 0,1135 | -2,00 | 0,050 | t |
| Homozygous RLL | Intercept | -7,7914 | 0,4447 | -17,52 | <0,001 | t |
| | Population | 0,1770 | 0,1440 | 1,23 | 0,224 | t |
| | Site | 0,7431 | 0,1563 | 4,76 | **<0,001** | t |
| | Coverage | 0,0655 | 0,0248 | 2,64 | **0,010** | t |
| | Batch | 0,2799 | 0,1801 | 1,55 | 0,125 | t |

**Table S8.** Comparison of genetic load between Guanapo (source population, in the intercept) and upper Turure (translocated population) including coverage as a covarate. Reported statistics are t-values for linear models and z-values for generalized linear models. Significant P-values are shown in bold.

| Dependent | Term | Estimate | SE | Statistic | P | Stat_type |
|---|---|---|---|---|---|---|
| Total RGL | Intercept | 0,0105 | 0,0008 | 13,20 | <0,001 | t |
| | Population | -0,0030 | 0,0005 | -6,25 | **<0,001** | t |
| | Coverage | 0,0001 | 0,0001 | 1,41 | 0,171 | t |
| ln(Heterozygous RGL) | Intercept | -7,0426 | 0,2250 | -31,30 | <0,001 | t |
| | Population | -1,0022 | 0,1369 | -7,32 | **<0,001** | t |
| | Coverage | 0,0046 | 0,0211 | 0,22 | 0,831 | t |
| Homozygous RGL | Intercept | 0,0030 | 0,0013 | 2,31 | 0,029 | t |
| | Population | 0,0012 | 0,0008 | 1,51 | 0,144 | t |
| | Coverage | 0,0001 | 0,0001 | 0,45 | 0,655 | t |
| Total RML | Intercept | -1,5566 | 0,0578 | -26,95 | <0,001 | z |
| | Population | -0,1448 | 0,0382 | -3,79 | **<0,001** | z |
| | Coverage | 0,0135 | 0,0051 | 2,62 | **0,009** | z |
| Heterozygous RML | Intercept | -1,6355 | 0,0769 | -21,26 | <0,001 | z |
| | Population | -0,5107 | 0,0491 | -10,40 | **<0,001** | z |
| | Coverage | -0,0122 | 0,0071 | -1,72 | **0,085** | z |
| Homozygous RML | Intercept | -3,0407 | 0,0820 | -37,08 | <0,001 | z |
| | Population | 0,4183 | 0,0579 | 7,22 | **<0,001** | z |
| | Coverage | 0,0365 | 0,0068 | 5,37 | **<0,001** | z |
| Total RLL | Intercept | -5,4176 | 0,4395 | -12,33 | <0,001 | z |
| | Population | -0,9173 | 0,2748 | -3,34 | **0,001** | z |
| | Coverage | 0,0218 | 0,0407 | 0,54 | 0,592 | z |
| Heterozygous RLL | Intercept | -5,6920 | 0,5374 | -10,59 | <0,001 | z |
| | Population | -0,7619 | 0,3385 | -2,25 | **0,024** | z |
| | Coverage | 0,0042 | 0,0500 | 0,09 | 0,932 | z |
| Homozygous RLL | Intercept | -6,7871 | 0,7627 | -8,90 | <0,001 | z |
| | Population | -1,2232 | 0,4702 | -2,60 | **0,009** | z |
| | Coverage | 0,0569 | 0,0701 | 0,81 | 0,417 | z |

**Table S9**. Comparison of genetic load between Turure upper (in the intercept) and two downstream sites, including coverage and sequencing batch as covariates. Reported statistics are t for lms on RGL, z for glms for missense and loss of function. Significant P-values for the population effects are in bold.

| Dependent | Term | Estimate | SE | Statistic | P | Stat_type |
|---|---|---|---|---|---|---|
| Total RGL | Intercept | 0,0069 | 0,0008 | 8,82 | 0,000 | t |
| | Population TUR middle | -0,0040 | 0,0002 | -18,89 | **<0,001** | t |
| | Population TUR lower | -0,0027 | 0,0002 | -12,53 | **<0,001** | t |
| | Coverage | 0,0001 | 0,0001 | 2,62 | **0,012** | t |
| | Batch | 0,0006 | 0,0002 | 2,52 | **0,016** | t |
| Heterozygous RGL | Intercept | 0,0042 | 0,0006 | 7,17 | 0,000 | t |
| | Population TUR middle | -0,0011 | 0,0002 | -7,05 | **<0,001** | t |
| | Population TUR lower | -0,0003 | 0,0002 | -1,91 | 0,063 | t |
| | Coverage | 0,0000 | 0,0000 | -0,31 | 0,761 | t |
| | Batch | -0,0001 | 0,0002 | -0,76 | 0,453 | t |
| Homozygous RGL | Intercept | 0,0027 | 0,0009 | 3,14 | 0,003 | t |
| | Population TUR middle | -0,0029 | 0,0002 | -12,31 | **<0,001** | t |
| | Population TUR lower | -0,0024 | 0,0002 | -10,03 | **<0,001** | t |
| | Coverage | 0,0001 | 0,0001 | 2,57 | **0,014** | t |
| | Batch | 0,0007 | 0,0003 | 2,79 | **0,008** | t |
| Total RML | Intercept | -1,6668 | 0,0740 | -22,53 | 0,000 | z |
| | Population TUR middle | -0,4280 | 0,0257 | -16,65 | **<0,001** | z |
| | Population TUR lower | -0,1909 | 0,0202 | -9,43 | **<0,001** | z |
| | Coverage | 0,0107 | 0,0047 | 2,27 | **0,023** | z |
| | Batch | 0,0273 | 0,0237 | 1,15 | 0,250 | z |
| Heterozygous RML | Intercept | -2,0217 | 0,1022 | -19,78 | 0,000 | z |
| | Population TUR middle | -0,2527 | 0,0348 | -7,27 | **<0,001** | z |
| | Population TUR lower | -0,0311 | 0,0276 | -1,13 | 0,259 | z |
| | Coverage | -0,0187 | 0,0065 | -2,86 | **0,004** | z |
| | Batch | -0,1044 | 0,0340 | -3,07 | **0,002** | z |
| Homozygous RML | Intercept | -2,6597 | 0,0981 | -27,12 | 0,000 | z |
| | Population TUR middle | -0,5888 | 0,0355 | -16,58 | **<0,001** | z |
| | Population TUR lower | -0,3410 | 0,0274 | -12,44 | **<0,001** | z |
| | Coverage | 0,0365 | 0,0062 | 5,89 | **<0,001** | z |
| | Batch | 0,1358 | 0,0305 | 4,45 | **<0,001** | z |
| Total RLL | Intercept | -5,7008 | 0,5632 | -10,12 | 0,000 | z |
| | Population TUR middle | -0,0282 | 0,1929 | -0,15 | 0,884 | z |
| | Population TUR lower | 0,4329 | 0,1492 | 2,90 | **0,004** | z |
| | Coverage | -0,0199 | 0,0359 | -0,56 | 0,579 | z |
| | Batch | -0,0110 | 0,1942 | -0,06 | 0,955 | z |
| Heterozygous RLL | Intercept | -6,3963 | 0,7056 | -9,06 | 0,000 | z |
| | Population TUR middle | -0,4966 | 0,2807 | -1,77 | 0,077 | z |
| | Population TUR lower | 0,3836 | 0,1867 | 2,05 | **0,040** | z |
| | Coverage | -0,0014 | 0,0448 | -0,03 | 0,974 | z |
| | Batch | 0,1012 | 0,2448 | 0,41 | 0,679 | z |
| Homozygous RLL | Intercept | -6,2890 | 0,9322 | -6,75 | <0,001 | z |
| | Population TUR middle | 0,5470 | 0,2801 | 1,95 | 0,051 | z |
| | Population TUR lower | 0,5373 | 0,2501 | 2,15 | **0,032** | z |
| | Coverage | -0,0535 | 0,0599 | -0,89 | 0,372 | z |
| | Batch | -0,1972 | 0,3192 | -0,62 | 0,537 | z |
